# Controllable multi-halogenation of a non-native substrate by SyrB2 iron halogenase

**DOI:** 10.1101/2024.05.08.593161

**Authors:** R. Hunter Wilson, Sourav Chatterjee, Elizabeth R. Smithwick, Anoop R. Damodaran, Ambika Bhagi-Damodaran

**Affiliations:** Department of Chemistry, University of Minnesota, Twin Cities Minneapolis, MN, 55455, United States

**Keywords:** 2OG-dependent, C-H functionalization, halogenation, non-heme-iron, multi-halogenation

## Abstract

Geminal, multi-halogenated functional groups are widespread in natural products and pharmaceuticals, yet no synthetic methodologies exist that enable selective multi-halogenation of unactivated C-H bonds. Biocatalysts are powerful tools for late-stage C-H functionalization, as they operate with high degrees of regio-, chemo-, and stereoselectivity. 2-oxoglutarate (2OG)-dependent non-heme iron halogenases chlorinate and brominate aliphatic C-H bonds offering a solution for achieving these challenging transformations. Here, we describe the ability of a non-heme iron halogenase, SyrB2, to controllably halogenate non-native substrate alpha-aminobutyric acid (Aba) to yield mono-chlorinated, di-chlorinated, and tri-chlorinated products. These chemoselective outcomes are achieved by controlling the loading of 2OG cofactor and SyrB2 biocatalyst. By using a ferredoxin-based biological reductant for electron transfer to the catalytic center of SyrB2, we demonstrate order-of-magnitude enhancement in the yield of tri-chlorinated product that were previously inaccessible using any single halogenase enzyme. We also apply these strategies to broaden SyrB2’s reactivity scope to include multi-bromination and demonstrate chemoenzymatic conversion of the ethyl side chain in Aba to an ethylyne functional group. We show how steric hindrance induced by the successive addition of halogen atoms on Aba’s C_4_ carbon dictates the degree of multi-halogenation by hampering C_3_-C_4_ bond rotation within SyrB2’s catalytic pocket. Overall, our work showcases the synthetic potential of iron halogenases to facilitate multi-C-H functionalization chemistry.

## Introduction

Achieving direct and selective functionalization of aliphatic C-H bonds poses significant challenges in organic synthesis. Biocatalysis has emerged as a solution to this problem, as enzymes offer regio-, stereo-, and chemo-selective functional group installation in large yields.^1^ Non-heme iron enzymes, specifically those which require a 2-oxoglutarate (2OG) cofactor, are of specific interest due to their incredibly diverse reaction scope which includes epimerization^2^, hydroxylation^3,4^, desaturation^5–7^, epoxidation^8–10^, and aziridination^11^. Notably, 2OG-dependent non-heme iron halogenases (hereafter referred to as iron halogenases) can perform chlorination^12,13^, di-chlorination^14–20^, and bromination^14,16,21–23^ on unactivated C-H bonds of their native substrates, though all of these reactions compete with a hydroxylation side reaction. The most extensively researched class of iron halogenases are the ACP-dependent halogenases which require delivery of their substrates by covalent attachment to a phosphopantetheinyl (Ppt) group appended to an acyl-carrier protein (ACP).^12,13,15,19–22,24^ In recent years, free-standing halogenases have also been discovered that bind and halogenate substrates in their active sites without the need of an ACP.^16,25,26^ Iron halogenases have been identified to be responsible for the incorporation of mono-chlorinated, di-chlorinated, or tri-chlorinated functional groups in a variety of natural products (**Fig. 1a**). Many of these natural products possess antibiotic or antimicrobial properties, sparking substantial research interest in recent decades.^18,24,27–29^

**Figure 1.**
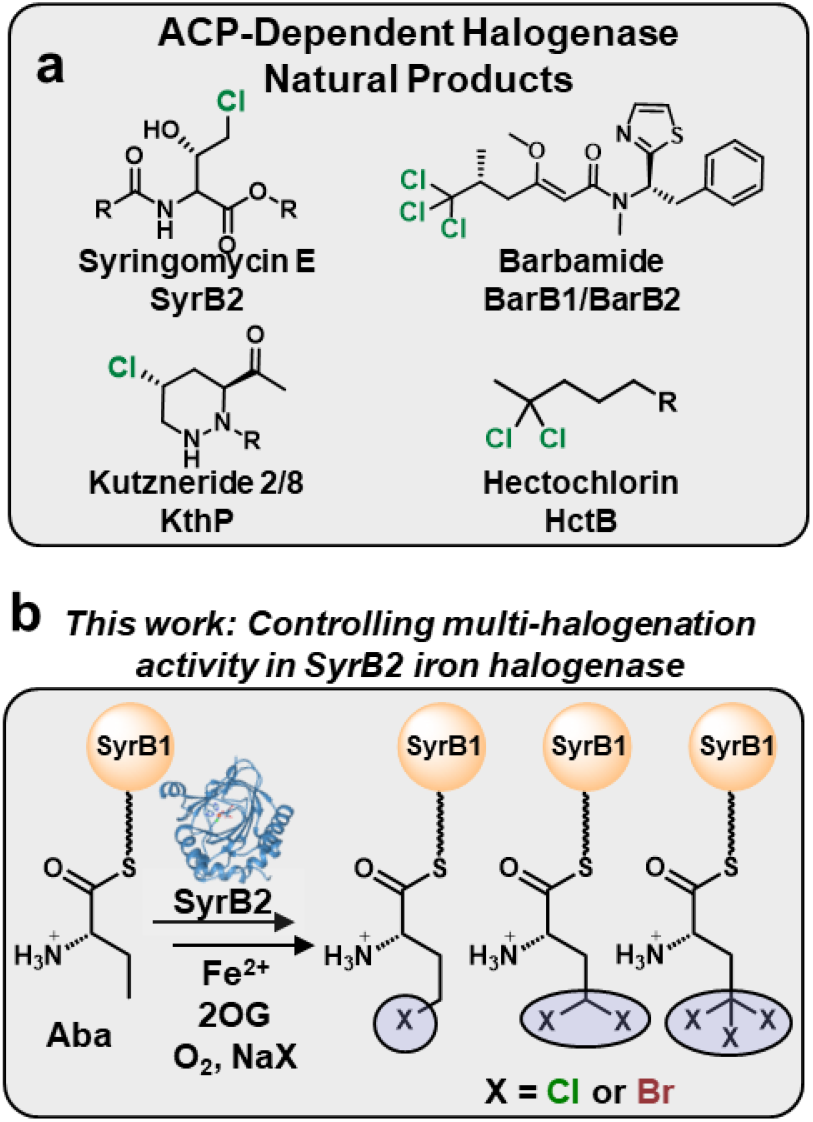
**a)** Halogenated natural products containing a variety of C-Cl_n_ bonds where n ranges from 1 to 3. Enzymes responsible for halogenated product formation are given under the compound name. **b)** This work demonstrates chemoselective control of C-H functionalization from mono- to tri-halogenation. This is achieved in a non-heme iron halogenase SyrB2 with SyrB1-Ppt-Aba substrate.

To date, no synthetic methodologies exist for direct multi-halogenation of unactivated C-H bonds.^30–35^ Existing methods for multi-halogenation almost always act upon pi bonds from alkenes/alkynes^36–45^, activated C-H bonds from ketones^46–48^, or require multi-step synthesis^49^ and few offer chemoselectivity towards the degree of halogenation^33,49–58^. In this work, we demonstrate systematic control of the degree of regioselective halogenation of a non-native substrate alpha-aminobutyric acid (Aba) by SyrB2 halogenase to yield mono-, di- and tri-chlorinated products. Using Molecular Dynamics (MD) simulations, we explore the molecular basis for the observed reactivity patterns and highlight the role of rotational flexibility of Aba within the SyrB2 catalytic pocket on the degree of halogenation. By using a ferredoxin-based biological reductant for electron transfer to the catalytic center of SyrB2, we demonstrate order-of-magnitude enhancement in the yield of tri-chlorinated product that were previously inaccessible using any single halogenase enzyme. We also reveal the ability of SyrB2 to perform di- and tri-bromination reactions and a potential application of geminal di-brominated Aba by converting it to an alkyne functional group useful for biorthogonal chemistry applications. Overall, our results offer new insights into controlling iron halogenase catalysis and expand their functionalization capabilities.

## Results and Discussion

### Controlling mono-vs. di-chlorination in SyrB2 halogenase using 2OG loading

We began our studies by investigating the reactivity of SyrB1-Ppt conjugated Aba (SyrB1-Ppt-Aba) with SyrB2 in the presence of varying equivalents of 2OG. The assay involves incubating SyrB2 with substrate SyrB1-Ppt-Aba and cofactors iron, 2OG, chloride and O_2_. The unreacted or functionalized Aba is subsequently released from SyrB1-Ppt-Aba via reaction with a thioesterase (TycF) and derivatized with 6-aminoquinolyl-n-hydroxysuccinimidyl carbamate (AQC) tag for UPLC separation and mass spectrometric analysis by Multiple Reaction Monitoring (MRM). As expected, the reaction of SyrB1-Ppt-Aba with SyrB2 in the absence of 2OG yielded no products underscoring the need of the 2OG cofactor to jumpstart catalysis via formation of a chloroferryl intermediate (**Fig. 2a, Figs. S1-3**). However, the addition of 0.5 equivalents of 2OG yielded <1% mono-chlorinated Aba (Cl-Aba) as a reaction product. We reasoned that the conversion to Cl-Aba could be low due to an intramolecular lactonization reaction, which has been demonstrated to be occurring in halogenated products with other non-heme iron enzymes (**Fig 2a**, inset).^14,16,59^ Subsequent UPLC/MRM studies validated such expectations with the major reaction product appearing in the MRM channel corresponding to the lactonized form of Aba (Lac-Aba), and was later confirmed via accurate mass measurements (**Fig. 2a, Figs. S4-6**). Moreover, by enhancing the 2OG cofactor to 1 equivalent and 2.5 equivalents with respect to substrate, we achieve 33% and 41% conversion to mono-chlorinated Aba product (Lac-Aba + Cl-Aba), respectively. Intriguingly, with these reaction conditions we also start to observe the formation of a product whose mass corresponds to di-chlorinated Aba (Cl_2_-Aba) (**Fig. S7**). We reasoned that the dichlorination of Aba could be happening by a second chlorination reaction on Aba’s C_4_ carbon. To validate this hypothesis and increase the conversion to di-chlorinated product, we increased the loading of 2OG cofactor. Indeed at 5 equivalents 2OG loading, we achieve a conversion of up to 97% Cl_2_-Aba, while decreasing the Lac-Aba and Cl-Aba to a combined 1.7% conversion (**Fig. 2a, Fig. S8**). Unlike Cl-Aba, Cl_2_-Aba appears to be resistant to intramolecular cyclization. We attribute this to the additional steric hinderance of a second chlorine atom on C_4_ that would prevent lactonization. As we further increased the 2OG loading to 10 and 100 eq., we also began to observe masses that correspond to a tri-chlorinated product (Cl_3_-Aba, up to 9% conversion) (**Figs. S8-S10**) that correlated with a decrease in conversion to Cl_2_-Aba (decreased to 87% conversion) (**Fig. 2a**). To confirm the identities of these analytes as Cl_2_-Aba and Cl_3_-Aba, we performed accurate mass measurements on our reaction products. These studies showed that the theoretical isotopic [M+H]^+^ distributions of Cl_2_-Aba and Cl_3_-Aba match those exhibited by the experimental isotopic distributions (**Fig. S11**). Moreover, the mass error between the theoretical and experimental monoisotopic masses are 2.3 ppm for Cl_2_-Aba and 1.6 ppm for Cl_3_-Aba, validating their identity. Overall, by systematic tuning of 2OG loading, we demonstrate controlled mono-vs. di-chlorination of Aba by SyrB2.

**Fig. 2:**
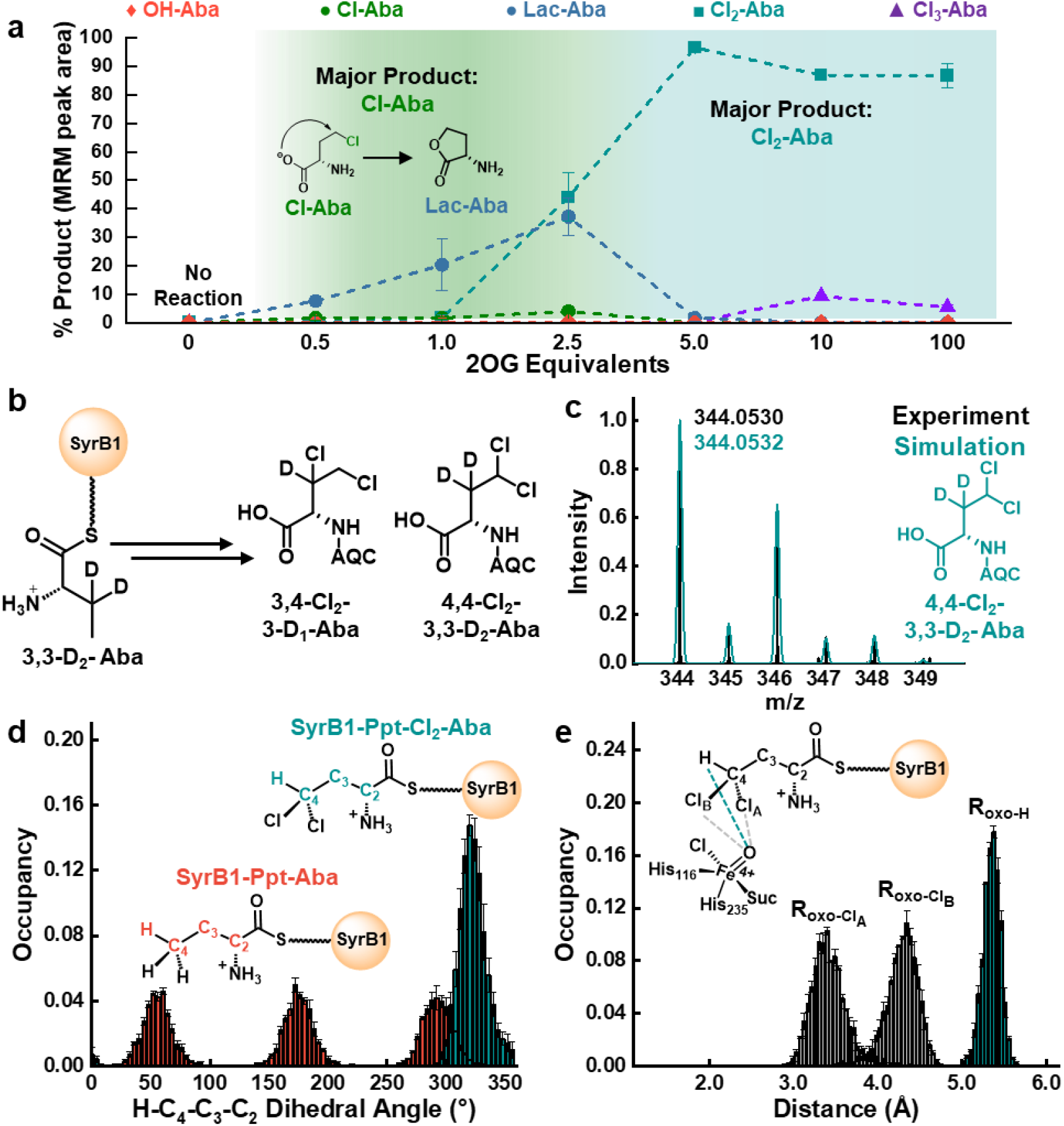
**a)** AQC-tagged products detected as a percentage of total MRM peak area for SyrB2’s reaction with SyrB1-Ppt-Aba at varied 2OG loading. Error bars are the standard deviation from n = 3 reactions. The % MRM peak area reflects trends in product selectivity and not definitive molar ratio. **b)** For a dichlorinated product, the chlorine atoms can both install on C_4_ or on C_3_ and C_4_. By conducting the reaction with a deuterated substrate probe, either both or a single deuterium can be retained on C_3_. **c)** Experimental HR-MS spectrum of the Cl_2_-Aba product from the chlorination reaction of 3,3-D_2_-Aba overlaid with the simulated spectrum for 4,4,Cl_2_-3,3-D_2_-Aba. **d)** MD simulation results monitoring the H-C_4_-C_3_-C_2_ dihedral angle of the SyrB1-Ppt-Aba and SyrB1-Ppt-Cl_2_-Aba substrates in the SyrB2 active site. **e)** MD simulation results monitoring the C_4_ chlorine and hydrogen atom distances of SyrB1-Ppt-Cl_2_-Aba to the oxo ligand of the chloroferryl intermediate in the SyrB2 active site. MD error bars are the standard error across three independent production runs. Suc = succinate.

While our accurate mass studies confirmed the identities of Cl_2_-Aba and Cl_3_-Aba, we wanted to structurally characterize the reaction products. More specifically, we wanted to confirm that the C_4_ position was the site where all new C-Cl bonds were formed. Given the small scale of our reactions and our existing detection by MS, we opted to use a deuterated substrate to probe the site-selectivity of the reaction. By appending 3,3-D_2_-Aba to SyrB1-Ppt and monitoring the chlorination assay products via MS, we would be able to detect how many deuterium atoms were retained in the Cl_2_-Aba product (**Fig. 2b**). If the Cl_2_-Aba product only contained a single deuterium atom, then we could infer that one chlorine atom was incorporated at C_3_ while the other was incorporated at C_4_. If both deuterium atoms were retained, then all C-Cl bonds would be formed at position C_4_, forming a geminal di-chlorinated product. In turn, we performed the chlorination assay with SyrB1-Ppt-3,3-D_2_-Aba substrate and probed the identity of Cl_2_-products using accurate mass measurements. When comparing our experimental result for the di-chlorinated product with 3,4,Cl_2_-3-D_1_-Aba, we see a poor fit to the isotopic distribution pattern and a monoisotopic mass error of 2933 ppm (**Fig. S12**). On the other hand, when comparing with the simulated mass spectrum in which both deuterium atoms are retained (4,4-Cl_2_-3,3-D_2_-Aba), we observe excellent agreement with the isotopic distribution pattern and a monoisotopic mass error of 0.6 ppm (**Fig. 2c**). These observations support that both deuterium atoms were retained and that all chlorine atoms were installed onto position C_4_. Next, we wanted to confirm that C_4_ was the position at which all three chlorine atoms were being installed for the Cl_3_-Aba product as well. Upon comparing the experimental mass spectrum to the simulated spectrum of 3,4,4-Cl_3_-3-D_1_-Aba, a poor fitting to the isotopic pattern is observed, and the monoisotopic mass error is 2671 ppm (**Fig. S13**). When overlaying the experimental spectrum and the simulated spectrum for 4,4,4-Cl_3_-3,3-D_2_-Aba, we see an excellent agreement between the experimental and simulated isotopic patterns and a monoisotopic mass error of 1.6 ppm (**Fig. S13**). This result shows that both deuterium atoms were retained during the assay and all three chlorine atoms were installed at the C_4_ position.

Overall, these studies demonstrate that SyrB2 can regioselectively multi-chlorinate Aba to produce geminal di- and tri-chlorinated products.

### Influence of substrate C-C bond rotational flexibility on the degree of chlorination

Our results so far demonstrate that chlorination reactions with equimolar SyrB1-Ppt-Aba and SyrB2 in the presence of excess 2OG yields di-chlorinated Aba with high selectivity (**Fig. 2a**). DFT bond dissociation energy (BDE) calculations reveal that C-H bond energy at the C_4_ position of Aba is lowered upon successive chlorination reactions suggesting that bond strength is not a factor for hindered multi-chlorination (**Fig. S14**). To understand why a third chlorination reaction at the C_4_ position is disfavored, we investigated the molecular basis for the degree of chlorination. For a chlorination reaction to proceed, the hydrogen atoms on Aba’s C_4_ must approach distances proximal to the oxo ligand of the reactive chloroferryl intermediate so it can be abstracted (**Fig. S3**). We hypothesized that the addition of two chlorine atoms to the C_4_ of Aba would introduce significant steric bulk and alter the conformational positioning of the substrate in the active site, disfavoring the third H-atom abstraction needed to form Cl_3_-Aba. We explored this hypothesis by performing MD simulations of SyrB2 with both the SyrB1-Ppt-Aba and SyrB1-Ppt-Cl_2_-Aba substrates. First, we compared the rotational flexibility of the C_3_-C_4_ bond between the two substrates by observing the H-C_4_-C_3_-C_2_ dihedral angle (**Fig. 2d**). For SyrB1-Ppt-Aba, we observe three distinct dihedral distributions, indicating facile rotation about the C_3_-C_4_ bond in SyrB2 active site. When comparing the distance distributions between the SyrB1-Ppt-Aba C_4_ hydrogens and the oxo ligand, we observe that they all overlay and can approach distances within <3.0 Å of the oxo ligand (**Fig. S15**). This implies that any of the hydrogens of SyrB1-Ppt-Aba can be easily abstracted to form a mono-chlorinated product. In contrast to the non-functionalized Aba substrate, SyrB1-Ppt-Cl_2_-Aba only has a single dihedral distribution, indicating that the C_3_-C_4_ bond isn’t rotatable, and the substrate is locked into a single dihedral conformation (**Fig 2d**). As a result, when we monitor the distance distribution between the C_4_’s chlorine and hydrogen atoms to the oxo ligand, each has a distinct distribution (Fig. 2e). While the chlorine atoms are more proximal to the oxo ligand (4.5 Å or less; shaded gray), the last abstractable hydrogen atom is located ∼5.5 Å from the oxo ligand (shaded cyan). At these longer distances, H-atom abstraction would be greatly impeded,^60^ which could explain hindered production of Cl_3_-Aba in our assays. In turn, the MD simulations suggest that steric effects upon successive chlorination can impact the optimal positioning of the substrate’s C_4_ hydrogen and may require multiple binding events thereby hindering tri-chlorination. Overall, our studies reveal that the steric bulk caused by the subsequent addition of chlorine atoms on the substrate can severely impede C_3_-C_4_ bond rotation in the SyrB2 active site and dictates the degree of multi-chlorination reactivity.

### Enhanced geminal tri-chlorinated product yields with ferredoxin as a biological reductant

The ability to obtain ∼9% Cl_3_-Aba in our halogenase assays was an exciting reaction outcome as no sole enzyme has been shown to directly tri-chlorinate a substrate, let alone one with unactivated, aliphatic C-H bonds. Notably, the biosynthesis of geminal tri-chlorinated functional group of barbamide requires the action of two independent iron halogenases, BarB1 and BarB2.^20^ More specifically, while BarB2 di-chlorinates the unactivated C_5_ carbon of leucine substrate (attached to BarA-Ppt ACP) (**Fig. 1a**), BarB1 completes the third chlorination step to yield the final tri-chlorinated product. In an effort to increase the yield of Cl_3_-Aba in our assays, we systematically varied the catalyst : substrate ratio (SyrB2 : SyrB1-Ppt-Aba) (**Fig. 3a**). When the substrate is in excess of the catalyst (SyrB2 : SyrB1-Ppt-Aba ratios of 1:4 and 1:2), we observe Cl_2_-Aba as the primary product with >98% conversion revealing how selective the enzyme is towards dichlorination. At equimolar catalyst : substrate ratio, however, we start to observe the appearance of ∼5% Cl_3_-Aba and the conversion of Cl_2_-Aba decreases to 87%. When we overload the amount of catalyst with respect to substrate (SyrB2 : SyrB1-Ppt-Aba ratios of 2:1 and 4:1), we observe significant amounts of tri-chlorinated product. Notably, at a catalyst : substrate ratio of 4:1, we observe as high as 72% of Aba converted to Cl_3_-Aba, with only 25% converted to Cl_2_-Aba (**Fig. 3a**). We believe that the need for excess catalyst-complex for tri-chlorination reaction arises from the limited turnover number of 7 ± 2 for SyrB2 before inactivation.^12^ MD simulations have shown that the di-chlorinated substrate, SyrB1-Ppt-Cl_2_-Aba, is locked in a dihedral conformation which can impede the third H-atom abstraction (**Fig 2d**,**e**). In turn, SyrB1-Ppt-Cl_2_-Aba bound to SyrB2 in an unfavorable locked single-dihedral conformation will need to disassociate from its partner and subsequently bind to an active SyrB2 in a dihedral conformation that is favorable for tri-chlorination. The opportunity for this to occur is significantly enhanced in the presence of excess catalyst. We reasoned that the low turnover for SyrB2 is likely related to inactivation via Fe^2+^ oxidation, a common uncoupling pathway in 2OG-utilizing iron enzymes.^61–64^ Such inactivation can be countered using a reductant such as ascorbate to reduce oxidized iron. However, ascorbate can also perform a nucleophilic attack on the thioester moiety of SyrB1-Ppt-Aba to release the tethered amino acid from the Ppt arm. Consequently, all assays with ACP-dependent halogenases till date have been conducted without a reductant. To address this challenge, we used a biological ferredoxin-based reductant system. With a significantly low redox potential of −410 mV^65^, ferredoxin is anticipated to reduce inactivated SyrB2 and bring it back in the catalytic cycle. Indeed, in the presence of ferredoxin, we observe an 11-fold enhancement in Cl_3_-Aba yields with as high as 61% of Aba converted to Cl_3_-Aba and with 31% converted to Cl_2_-Aba at a catalyst : substrate ratio of 1:1 (**Fig. 3b**). Overall, these results along with our computational findings, help understand what factors enable or limit tri-chlorination reactions in iron halogenases.

**Fig. 3:**
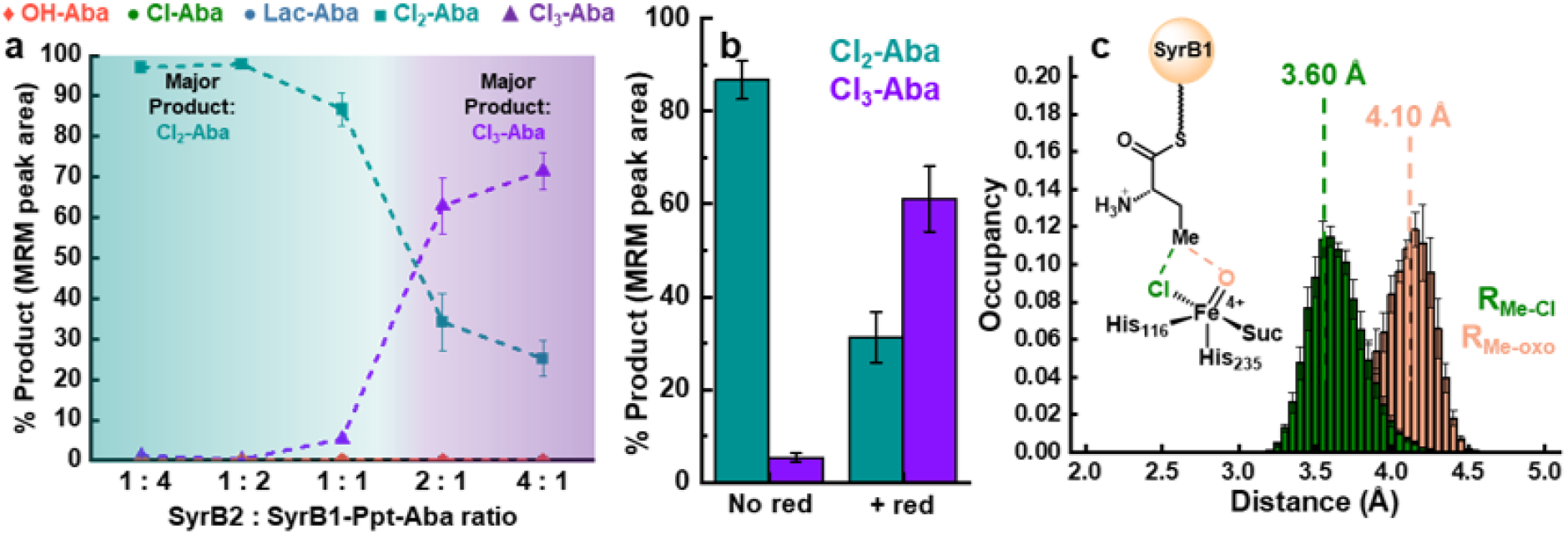
AQC-tagged products detected as a percentage of total MRM peak area for SyrB2’s reaction with SyrB1-Ppt-Aba at varied SyrB2 biocatalyst loading. No AQC-tagged mono-chlorinated, lactonized, or hydroxylated Aba analytes were detected in the assays. The 2OG loading was set to 100 equivalents; error bars are the standard deviation from n = 3 reactions. **b)** AQC-tagged products detected as a percentage of total MRM peak area for SyrB2’s reaction with SyrB1-Ppt-Aba without and with the biological reductant (abbreviated as red). The substrate to catalyst loading ratio was kept at 1:1 and the 2OG loading was set to 100 equivalents. Error bars are the standard deviation from n = 3 reactions. The % MRM peak area in a-b reflects trends in product selectivity and not definitive molar ratio. **c)** MD simulations of the SyrB2/SyrB1-Ppt-Aba complex with the reactive chloroferryl intermediate show that the reactive carbon is positioned closer to the halide ligand, favoring a halogenation outcome; error bars are the standard error over three independent production runs and vertical lines indicate the distance at which 50% of the distribution occurs.

### Molecular basis for chlorination over hydroxylation with non-native Aba substrate

To establish a molecular basis for our chemoselective chlorination outcome, we conducted MD simulations of the SyrB2/SyrB1-Ppt-Aba complex. We modelled the reactive chloroferryl intermediate in the SyrB2 active site as a distorted five-coordinate chloroferryl isomer with the oxo and chloro ligands in the equatorial plane as spectroscopically identified previously and monitored the distances between the reactive carbon center (C_4_) and the oxo and chloro ligands.^63^ Halogenases tend to organize the reactive carbon of their substrates closer to the halide ligand to favor rebound with the halogen upon H-atom abstraction by the haloferryl intermediate (**Fig. S3**).^66–72^ We observe that the reactive methyl group of Aba orients itself 0.5 Å closer to the chloride ligand than the oxo ligand over 1.5 μs of simulations (**Fig. 3c**). This preference in distance distribution suggests that SyrB1-Ppt-Aba is positioned for a halogenation outcome over hydroxylation, which is validated by our experimental results (**Figs. 2a, 3a**). In our studies, the MRM channels corresponding to OH-Aba show negligible peaks with no difference between the reactions of SyrB1-Ppt-Aba with SyrB2 or without SyrB2, indicating chemoselective chlorination of Aba by SyrB2. We note that we have employed a thioesterase, TycF, for liberating Aba which is a well-established method for releasing amino acids tethered to Ppt arm across various ACPs.^13,14,19,73–76^ Previous studies that suggested similar propensity of Aba to halogenate and hydroxylate did not discuss the potential for cyclization of Cl-Aba or the potential for other side products (including OH-Aba) from the basic conditions of hydroxylamine derivatization (**Fig. S16**)._60,77_

### Unprecedented multi-bromination activity

Thus far, we’ve demonstrated control over the degree of multi-chlorination, however, iron halogenases are also capable of mono-brominating their native substrates. ^14,16,23,59,71^ Given the propensity for the SyrB1-Ppt-Aba substrate to form multi-chlorinated products, next, we investigated if this reactivity could be extended to multi-bromination. We note that multi-bromination reactions are harder to accomplish as H-abstraction, ferryl decay and radical rebound are all rendered slower for bromination as compared to chlorination.^22,78,79^ As such, there are no previous reports of di- and tri-bromination by iron halogenases. To characterize bromination activity, we applied methods for controlling selectivity that were described above for multi-chlorination. First, we tested how the equivalents of 2OG cofactor affected the percent conversion for all products (**Fig. 4a, Figs. S17-23**). In the absence of 2OG, no products were observed, indicating that the addition of 2OG is required to initiate catalysis. Upon increasing the addition of 2OG up to 2.5 equivalents, the Lac-Aba product becomes the dominant product. Akin to the chlorination reaction, Lac-Aba forms from an intramolecular S_N_2 reaction in which the carboxylate group attacks the C-Br bond (**Fig. 4a**, inset). Unlike the chlorination reactions, however, no mono-brominated product (Br-Aba) was observed likely due to bromide being a better leaving group than chloride for the lactonization reaction. As we increased the 2OG loading to 5 equivalents, a product with masses corresponding to di-brominated Aba (Br_2_-Aba) became the major product, with a still appreciable amount of Lac-Aba present. In our chlorination assays, the Cl_2_-Aba product became the majority product after adding 2.5 equivalents of 2OG and was the dominant product after adding 5 equivalents (**Fig. 2**). Since bromination isn’t the native reaction for SyrB2, it appears that bromination operates slower than chlorination as seen in kinetics assays with iron halogenases^21,22,78,79^, leading to a lower percent conversion to multi-brominated products. By adding 10 eq. of 2OG, we’re able to increase the conversion of Br_2_-Aba up to 85%, concomitant with a decrease in Lac-Aba (and thereby Br-Aba). At 10 equivalents 2OG, a product with masses consistent with a tri-brominated product (Br_3_-Aba) begins to appear. By increasing the 2OG loading to 100 equivalents, we increase the conversion of Br_3_-Aba up to 10%. Overloading 2OG at this amount also eliminates signals associated with Lac-Aba, thereby fully shifting the chemoselectivity to multi-brominated Aba. Analogous to our chlorination assays, negligible OH-Aba was detected at any amount of 2OG loading during these assays. Overall, our studies reveal the ability to modulate chemoselectivity between mono-brominated and multi-brominated products by altering the levels of 2OG cofactor present in the halogenation reaction.

**Fig. 4:**
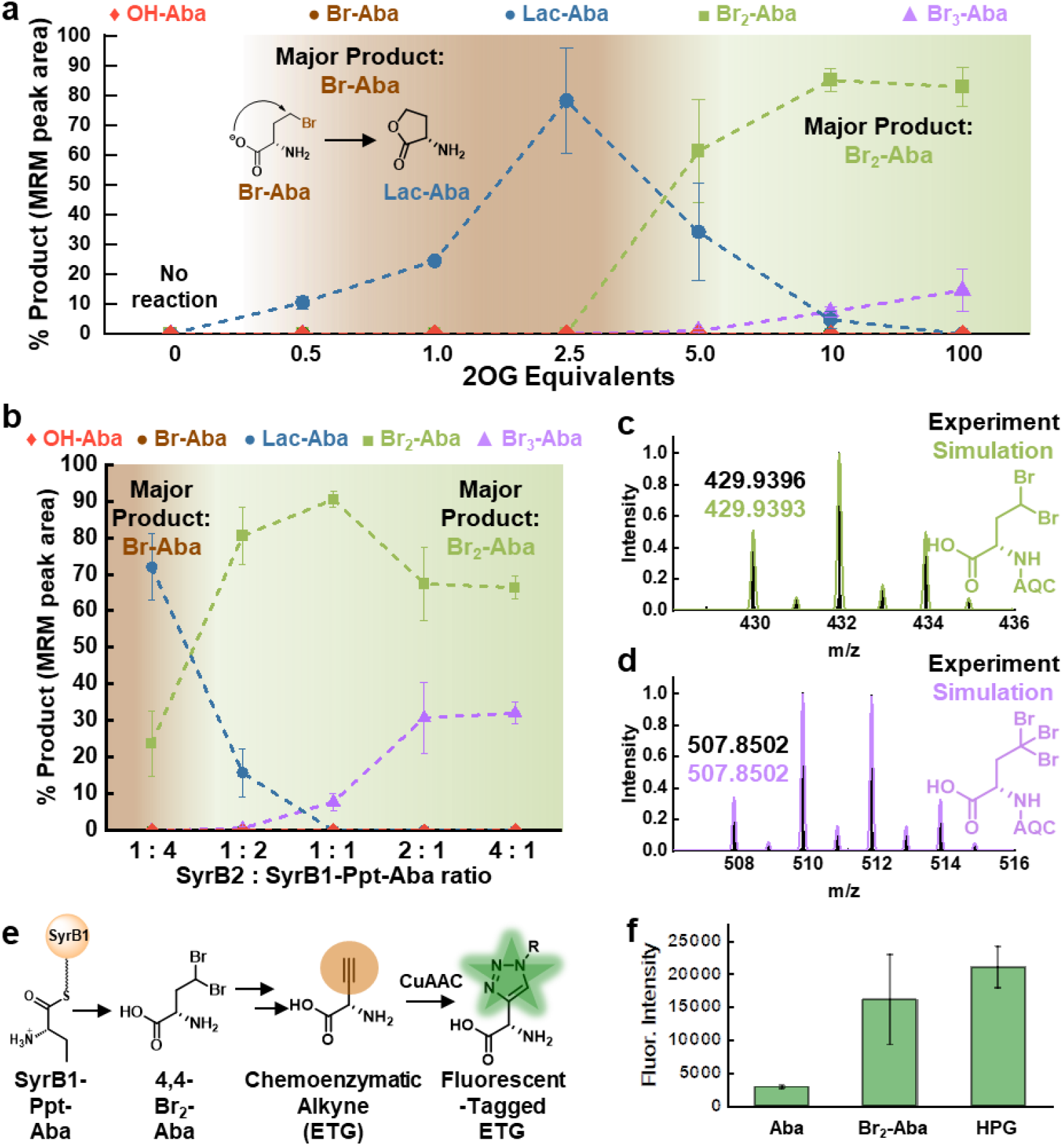
Iron halogenase catalysed multi-bromination activity and demonstrated synthetic utility. AQC-tagged products detected as a percentage of total MRM peak area for SyrB2’s reaction with SyrB1-Ppt-Aba at varied **a)** 2OG loading and **b**) varied SyrB2 biocatalyst loading. Error bars are the standard deviation from n = 3 reactions. The % MRM peak area in a-b reflects trends in product selectivity and not definitive molar ratio. **c)** HR-MS spectrum validating Br_2_-Aba identity. **d)** HR-MS spectrum validating Br_3_-Aba identity. **e)** Overview for the chemoenzymatic conversion of Aba to ethynylglycine (ETG) and its detection via click chemistry assay. **f)** Fluorescence responses demonstrating chemoenzymatic alkyne formation from Br_2_-Aba and a 10 μM homopropargylglycine (HPG) standard. Error bars are the standard deviation from three independent reactions/assays.

In an attempt to increase the conversion to Br_3_-Aba, we again varied the biocatalyst loading by altering the ratio of SyrB2 : SyrB1-Ppt-Aba (**Fig. 4b**). At a catalyst : substrate ratio of 1:4, we observe Lac-Aba as the dominant product at 72% conversion while Br_2_-Aba is the minor product at 24% conversion. This contrasts with our chlorination studies that revealed Cl_2_-Aba as the major product (>95%) at 1:4 catalyst : substrate ratio suggesting the lower potency for multi-turnovers with bromide as compared to chloride. However, as we increase the catalyst loading for bromination with catalyst : substrate ratios of 1:2 and 1:1, we start observing Br_2_-Aba as the major product with as high as 90% conversions. Finally, when catalyst is in excess of the substrate with catalyst : substrate ratios of 2:1 and 4:1, we increase the conversion of Br_3_-Aba up to 32% which is notable but considerably less than 72% Cl_3_-Aba observed for chlorination reactions under similar conditions. Unlike tri-chlorination, incorporating a ferredoxin-based biological reductant in our tri-bromination assay did not enhance the yield of tri-brominated product, reaffirming the difficulty of accomplishing bromination and suggesting the role of redox potential effects (**Fig. S24**). To verify our multi-brominated analytes’ identity, we obtained accurate mass spectra of these products. For the Br_2_-Aba species, we observe a match between the theoretical and experimental isotopic distribution patterns and a monoisotopic mass error of 0.7 ppm (**Fig. 4c**). Comparison between the experimental and theoretical Br_3_-Aba accurate mass spectra gives a good agreement for the isotopic distribution pattern and a monoisotopic mass error of 0.0 ppm (**Fig. 4d**). Moreover, we confirmed C_4_ of Aba as the site of functionalization for multi-bromination using the deuterium labelled SyrB1-Ppt-3,3-D_2_-Aba substrate similar to our chlorination studies (**Figs. S25, 26**). Our studies suggest that for both Br_2_-Aba and Br_3_-Aba, all bromine atoms were installed at C_4_ and the regioselective outcome of C-H functionalization does not differ between the native chlorination and non-native bromination reactions. To highlight the utility of such multi-brominated amino acid products, we performed dehydrohalogenation of SyrB2 generated Br_2_-Aba to ethynylglycine (ETG), an alkyne analog useful for click chemistry reactions and easily detected via an azide– alkyne cycloaddition (CuAAc) click reaction with CalFluor 488 Azide (**Fig. 4e, f**). Ultimately, these studies mark the first instance of an iron halogenase exhibiting multi-bromination activity which further broadens their already diverse catalytic repertoire.

## Conclusion

The ability of iron halogenases to chlorinate unactivated C(*sp*^*3*^)-H bonds with high chemo- and regioselectivity is highly desirable from a chemical synthesis perspective. We demonstrate how the degree of halogenation can be controlled in SyrB2 halogenases to obtain chemo- and regioselective multi-halogenation of Aba, likely due to the proximal positioning of C_4_ towards the chloro ligand over the oxo ligand of the reactive chloroferryl intermediate. Our studies reveal unprecedented tri-chlorination (aided by use of a biological reductant system) and multi-bromination activity in SyrB2, expanding the catalytic prowess of iron halogenases. From a mechanistic perspective, we discuss the importance of the substrate’s C-C bond rotational flexibility at the site of C-H functionalization in controlling the degree of halogenation. From a chemical synthesis perspective, our findings reveal the utility of iron halogenases towards late-stage multi-C-H functionalization. Insights gained from our work on controlling the degree of C-H functionalization could find use in other C-H functionalizing metalloenzymes as well.^80–83^ Overall, molecular insights gained from our work have the potential to guide the design of next-generation high performance multi-halogenation biocatalysts.

## Supporting information

Supp Info_multihalogenation

## ASSOCIATED CONTENT

### Supporting Information

Materials and Methods, UPLC-MRM chromatograms, reaction cycle of 2OG-dependent non-heme iron halogenases, accurate mass spectra of analytes, multi-halogenation reactivity of Thr, Aba, and Val as a function of 2OG equivalents, Bond-dissociation energy DFT calculated energies, Distance distribution of Aba C_4_ Hydrogens to haloferryl intermediate, possible hydroxamate reaction products, multi-bromination analysis with Fdx/FdxR reduction system, additional references

## AUTHOR INFORMATION

## Author Contributions

The manuscript was written through contributions of all authors. All authors have given approval to the final version of the manuscript.

## ACKNOWLEDGMENT

ERS and RHW acknowledge the support of the National Institute of Health Chemical Biology Training Grant (T32GM132029). This work was supported by NSF CBET (Grant # 2046527). Mass spectrometry analysis was performed at The University of Minnesota Department of Chemistry Mass Spectrometry Laboratory (MSL), supported by the Office of the Vice President of Research, College of Science and Engineering, and the Department of Chemistry at the University of Minnesota. The content of this paper is the sole responsibility of the authors and does not represent endorsement by MSL personnel. The authors would like to thank Profs. Krebs and Bollinger (Penn State) for providing SyrB1, SyrB2, and Sfp plasmids and Prof. Balskus (MIT) for providing TycF plasmid.

## ABBREVIATIONS

2OG: 2-oxoglutarate
ACP: acyl-carrier protein
Ppt: phosphopantetheine
Aba: alpha-aminobutyric acid
MD: Molecular Dynamics
AQC: 6-aminoquinolyl-n-hydroxysuccinimidyl carbamate
MRM: Multiple Reaction Monitoring

## REFERENCES

(1) Bell, E. L.; Finnigan, W.; France, S. P.; Green, A. P.; Hayes, M. A.; Hepworth, L. J.; Lovelock, S. L.; Niikura, H.; Osuna, S.; Romero, E.; Ryan, K. S.; Turner, N. J.; Flitsch, S. L. Biocatalysis. Nat Rev Methods Primers 2021, 1 (1), 1–21. 10.1038/s43586-021-00044-z.

(2) Chang, W.; Guo, Y.; Wang, C.; Butch, S. E.; Rosenzweig, A. C.; Boal, A. K.; Krebs, C.; Bollinger, J. M. Mechanism of the C5 Stereoinversion Reaction in the Biosynthesis of Carbapenem Antibiotics. Science 2014, 343 (6175), 1140–1144. 10.1126/science.1248000.

(3) McDonough, M. A.; Li, V.; Flashman, E.; Chowdhury, R.; Mohr, C.; Liénard, B. M. R.; Zondlo, J.; Oldham, N. J.; Clifton, I. J.; Lewis, J.; McNeill, L. A.; Kurzeja, R. J. M.; Hewitson, K. S.; Yang, E.; Jordan, S.; Syed, R. S.; Schofield, C. J. Cellular Oxygen Sensing: Crystal Structure of Hypoxia-Inducible Factor Prolyl Hydroxylase (PHD2). Proceedings of the National Academy of Sciences 2006, 103 (26), 9814–9819. 10.1073/pnas.0601283103.

(4) Bollinger Jr., J. M.; Price, J. C.; Hoffart, L. M.; Barr, E. W.; Krebs, C. Mechanism of Taurine: α-Ketoglutarate Dioxygenase (TauD) from Escherichia Coli. European Journal of Inorganic Chemistry 2005, 2005 (21), 4245–4254. 10.1002/ejic.200500476.

(5) Dunham, N. P.; Mitchell, A. J.; Del Río Pantoja, J. M.; Krebs, C.; Bollinger, J. M. Jr.; Boal, K. α-Amine Desaturation of d-Arginine by the Iron(II)- and 2-(Oxo)Glutarate-Dependent l-Arginine 3-Hydroxylase, VioC. Biochemistry 2018, 57 (46), 6479–6488. 10.1021/acs.biochem.8b00901.

(6) Dunham, N. P.; Chang, W.; Mitchell, A. J.; Martinie, R. J.; Zhang, B.; Bergman, J. A.; Rajakovich, L. J.; Wang, B.; Silakov, A.; Krebs, C.; Boal, A. K.; Bollinger, J. M. Jr. Two Distinct Mechanisms for C–C Desaturation by Iron(II)- and 2-(Oxo)Glutarate-Dependent Oxygenases: Importance of α-Heteroatom Assistance. J. Am. Chem. Soc. 2018, 140 (23), 7116–7126. 10.1021/jacs.8b01933.

(7) Kim, W.; Chen, T.-Y.; Cha, L.; Zhou, G.; Xing, K.; Canty, N. K.; Zhang, Y.; Chang, W. Elucidation of Divergent Desaturation Pathways in the Formation of Vinyl Isonitrile and Isocyanoacrylate. Nat Commun 2022, 13 (1), 5343. 10.1038/s41467-022-32870-4.

(8) Liu, P.; Mehn, M. P.; Yan, F.; Zhao, Z.; Que, Lawrence; Liu, H. Oxygenase Activity in the Self-Hydroxylation of (S)-2-Hydroxypropylphosphonic Acid Epoxidase Involved in Fosfomycin Biosynthesis. J. Am. Chem. Soc. 2004, 126 (33), 10306–10312. 10.1021/ja0475050.

(9) Li, J.; Liao, H.-J.; Tang, Y.; Huang, J.-L.; Cha, L.; Lin, T.-S.; Lee, J. L.; Kurnikov, I. V.; Kurnikova, M. G.; Chang, W.; Chan, N.-L.; Guo, Y. Epoxidation Catalyzed by the Nonheme Iron(II)- and 2-Oxoglutarate-Dependent Oxygenase, AsqJ: Mechanistic Elucidation of Oxygen Atom Transfer by a Ferryl Intermediate. J. Am. Chem. Soc. 2020, 142 (13), 6268–6284. 10.1021/jacs.0c00484.

(10) Busby, R. W.; Townsend, C. A. A Single Monomeric Iron Center in Clavaminate Synthase Catalyzes Three Nonsuccessive Oxidative Transformations. Bioorganic & Medicinal Chemistry 1996, 4 (7), 1059–1064. 10.1016/0968-0896(96)00088-0.

(11) Cha, L.; Paris, J. C.; Zanella, B.; Spletzer, M.; Yao, A.; Guo, Y.; Chang, W. Mechanistic Studies of Aziridine Formation Catalyzed by Mononuclear Non-Heme Iron Enzymes. J. Am. Chem. Soc. 2023, 145 (11), 6240–6246. 10.1021/jacs.2c12664.

(12) Vaillancourt, F. H.; Yin, J.; Walsh, C. T. SyrB2 in Syringomycin E Biosynthesis Is a Nonheme FeII α-Ketoglutarate- and O2-Dependent Halogenase. PNAS 2005, 102 (29), 10111–10116. 10.1073/pnas.0504412102.

(13) Jiang, W.; Heemstra, J. R. Jr.; Forseth, R. R.; Neumann, C. S.; Manaviazar, S.; Schroeder, F. C.; Hale, K. J.; Walsh, C. T. Biosynthetic Chlorination of the Piperazate Residue in Kutzneride Biosynthesis by KthP. Biochemistry 2011, 50 (27), 6063–6072. 10.1021/bi200656k.

(14) Vaillancourt, F. H.; Vosburg, D. A.; Walsh, C. T. Dichlorination and Bromination of a Threonyl-S-Carrier Protein by the Non-Heme FeII Halogenase SyrB2. ChemBioChem 2006, 7 (5), 748–752. 10.1002/cbic.200500480.

(15) Ueki, M.; Galonić, D. P.; Vaillancourt, F. H.; Garneau-Tsodikova, S.; Yeh, E.; Vosburg, D. A.; Schroeder, F. C.; Osada, H.; Walsh, C. T. Enzymatic Generation of the Antimetabolite γ,γ-Dichloroaminobutyrate by NRPS and Mononuclear Iron Halogenase Action in a Streptomycete. Chemistry & Biology 2006, 13 (11), 1183–1191. 10.1016/j.chembiol.2006.09.012.

(16) Neugebauer, M. E.; Sumida, K. H.; Pelton, J. G.; McMurry, J. L.; Marchand, J. A.; Chang, M. C. Y. A Family of Radical Halogenases for the Engineering of Amino-Acid-Based Products. Nat Chem Biol 2019, 15 (10), 1009–1016. 10.1038/s41589-019-0355-x.

(17) Kissman, E. N.; Neugebauer, M. E.; Sumida, K. H.; Swenson, C. V.; Sambold, N. A.; Marchand, J. A.; Millar, D. C.; Chang, M. C. Y. Biocatalytic Control of Site-Selectivity and Chain Length-Selectivity in Radical Amino Acid Halogenases. Proceedings of the National Academy of Sciences 2023, 120 (12), e2214512120. 10.1073/pnas.2214512120.

(18) Ramaswamy, A. V.; Sorrels, C. M.; Gerwick, W. H. Cloning and Biochemical Characterization of the Hectochlorin Biosynthetic Gene Cluster from the Marine Cyanobacterium Lyngbya Majuscula. J. Nat. Prod. 2007, 70 (12), 1977–1986. 10.1021/np0704250.

(19) Pratter, S. M.; Ivkovic, J.; Birner-Gruenberger, R.; Breinbauer, R.; Zangger, K.; Straganz, G. D. More than Just a Halogenase: Modification of Fatty Acyl Moieties by a Trifunctional Metal Enzyme. ChemBioChem 2014, 15 (4), 567–574. 10.1002/cbic.201300345.

(20) Galonić, D. P.; Vaillancourt, F. H.; Walsh, C. T. Halogenation of Unactivated Carbon Centers in Natural Product Biosynthesis: Trichlorination of Leucine during Barbamide Biosynthesis. J. Am. Chem. Soc. 2006, 128 (12), 3900–3901. 10.1021/ja060151n.

(21) Galonić, D. P.; Barr, E. W.; Walsh, C. T.; Bollinger, J. M.; Krebs, C. Two Interconverting Fe(IV) Intermediates in Aliphatic Chlorination by the Halogenase CytC3. Nat Chem Biol 2007, 3 (2), 113–116. 10.1038/nchembio856.

(22) Galonić Fujimori, D.; Barr, E. W.; Matthews, M. L.; Koch, G. M.; Yonce, J. R.; Walsh, C. T.; Bollinger, J. M. Jr.; Krebs, C.; Riggs-Gelasco, P. J. Spectroscopic Evidence for a High-Spin Br-Fe(IV)-Oxo Intermediate in the α-Ketoglutarate-Dependent Halogenase CytC3 from Streptomyces. J. Am. Chem. Soc. 2007, 129 (44), 13408–13409. 10.1021/ja076454e.

(23) Zhu, Q.; Hillwig, M. L.; Doi, Y.; Liu, X. Aliphatic Halogenase Enables Late-Stage C−H Functionalization: Selective Synthesis of a Brominated Fischerindole Alkaloid with Enhanced Antibacterial Activity. ChemBioChem 2016, 17 (6), 466–470. 10.1002/cbic.201500674.

(24) Chang, Z.; Flatt, P.; Gerwick, W. H.; Nguyen, V.-A.; Willis, C. L.; Sherman, D. H. The Barbamide Biosynthetic Gene Cluster: A Novel Marine Cyanobacterial System of Mixed Polyketide Synthase (PKS)-Non-Ribosomal Peptide Synthetase (NRPS) Origin Involving an Unusual Trichloroleucyl Starter Unit. Gene 2002, 296 (1), 235–247. 10.1016/S0378-1119(02)00860-0.

(25) Hillwig, M. L.; Liu, X. A New Family of Iron-Dependent Halogenases Acts on Freestanding Substrates. Nat Chem Biol 2014, 10 (11), 921–923. 10.1038/nchembio.1625.

(26) Hillwig, M. L.; Zhu, Q.; Ittiamornkul, K.; Liu, X. Discovery of a Promiscuous Non-Heme Iron Halogenase in Ambiguine Alkaloid Biogenesis: Implication for an Evolvable Enzyme Family for Late-Stage Halogenation of Aliphatic Carbons in Small Molecules. Angewandte Chemie International Edition 2016, 55 (19), 5780–5784. 10.1002/anie.201601447.

(27) Orjala, J.; Gerwick, W. H. Barbamide, a Chlorinated Metabolite with Molluscicidal Activity from the Caribbean Cyanobacterium Lyngbya Majuscula. J. Nat. Prod. 1996, 59 (4), 427–430. 10.1021/np960085a.

(28) Ullrich, M.; Bender, C. L. The Biosynthetic Gene Cluster for Coronamic Acid, an Ethylcyclopropyl Amino Acid, Contains Genes Homologous to Amino Acid-Activating Enzymes and Thioesterases. J Bacteriol 1994, 176 (24), 7574–7586.

(29) Zhang, H.; Kakeya, H.; Osada, H. Biosynthesis of 1-Aminocyclopropane-1-Carboxylic Acid Moiety on Cytotrienin A in Streptomyces Sp. Tetrahedron Letters 1998, 39 (38), 6947–6948. 10.1016/S0040-4039(98)01460-9.

(30) Chung, W.; Vanderwal, C. D. Stereoselective Halogenation in Natural Product Synthesis. Angewandte Chemie International Edition 2016, 55 (14), 4396–4434. 10.1002/anie.201506388.

(31) Landry, M. L.; Burns, N. Z. Catalytic Enantioselective Dihalogenation in Total Synthesis. Acc. Chem. Res. 2018, 51 (5), 1260–1271. 10.1021/acs.accounts.8b00064.

(32) Dong, J.; Fernández-Fueyo, E.; Hollmann, F.; Paul, C. E.; Pesic, M.; Schmidt, S.; Wang, Y.; Younes, S.; Zhang, W. Biocatalytic Oxidation Reactions: A Chemist’s Perspective. Angewandte Chemie International Edition 2018, 57 (30), 9238–9261. 10.1002/anie.201800343.

(33) Das, R.; Kapur, M. Transition-Metal-Catalyzed Site-Selective C−H Halogenation Reactions. Asian Journal of Organic Chemistry 2018, 7 (8), 1524–1541. 10.1002/ajoc.201800142.

(34) Scheide, M. R.; Nicoleti, C. R.; Martins, G. M.; Braga, A. L. Electrohalogenation of Organic Compounds. Org. Biomol. Chem. 2021, 19 (12), 2578–2602. 10.1039/D0OB02459G.

(35) Nanjo, T.; Matsumoto, A.; Oshita, T.; Takemoto, Y. Synthesis of Chlorinated Oligopeptides via γ- and δ-Selective Hydrogen Atom Transfer Enabled by the N-Chloropeptide Strategy. J. Am. Chem. Soc. 2023. 10.1021/jacs.3c06931.

(36) Liang, L.; Guo, L.-D.; Tong, R. Achmatowicz Rearrangement-Inspired Development of Green Chemistry, Organic Methodology, and Total Synthesis of Natural Products. Acc. Chem. Res. 2022, 55 (16), 2326–2340. 10.1021/acs.accounts.2c00358.

(37) Fu, N.; Sauer, G. S.; Lin, S. Electrocatalytic Radical Dichlorination of Alkenes with Nucleophilic Chlorine Sources. J. Am. Chem. Soc. 2017, 139 (43), 15548–15553. 10.1021/jacs.7b09388.

(38) Sauer, G. S.; Lin, S. An Electrocatalytic Approach to the Radical Difunctionalization of Alkenes. ACS Catal. 2018, 8 (6), 5175–5187. 10.1021/acscatal.8b01069.

(39) Seidl, F. J.; Min, C.; Lopez, J. A.; Burns, N. Z. Catalytic Regio- and Enantioselective Haloazidation of Allylic Alcohols. J. Am. Chem. Soc. 2018, 140 (46), 15646–15650. 10.1021/jacs.8b10799.

(40) Lian, P.; Long, W.; Li, J.; Zheng, Y.; Wan, X. Visible-Light-Induced Vicinal Dichlorination of Alkenes through LMCT Excitation of CuCl2. Angewandte Chemie International Edition 2020, 59 (52), 23603–23608. 10.1002/anie.202010801.

(41) Sarie, J. C.; Neufeld, J.; Daniliuc, C. G.; Gilmour, R. Catalytic Vicinal Dichlorination of Unactivated Alkenes. ACS Catal. 2019, 9 (8), 7232–7237. 10.1021/acscatal.9b02313.

(42) Bai, Y.; Li, Y.; Zhang, Z.; Yang, X.; Zhang, J.; Chen, L.; Li, Y.; Zeng, X.; Zhang, M. Regioselective Difunctionalization of Alkene: A Simple Access to Haloether, Haloesters and Halohydrins. Tetrahedron Letters 2022, 101, 153923. 10.1016/j.tetlet.2022.153923.

(43) Cui, H.; Shen, Y.; Chen, Y.; Wang, R.; Wei, H.; Fu, P.; Lei, X.; Wang, H.; Bi, R.; Zhang, Y. Two-Stage Syntheses of Clionastatins A and B. J. Am. Chem. Soc. 2022, 144 (20), 8938–8944. 10.1021/jacs.2c03872.

(44) Jayaraman, A.; Cho, E.; Irudayanathan, F. M.; Kim, J.; Lee, S. Metal-Free Decarboxylative Trichlorination of Alkynyl Carboxylic Acids: Synthesis of Trichloromethyl Ketones. Advanced Synthesis & Catalysis 2018, 360 (1), 130–141. 10.1002/adsc.201701116.

(45) Li, X.; Jin, J.; Chen, P.; Liu, G. Catalytic Remote Hydrohalogenation of Internal Alkenes. Nat. Chem. 2022, 14 (4), 425–432. 10.1038/s41557-021-00869-x.

(46) Zhou, J.; Tang, D.; Bian, M. Facile Approach to Geminal Heterodihalogenation. One-Pot Synthesis of α-Bromo-α-Chloro Ketones. Synlett 2020, 31 (14), 1430–1434. 10.1055/s-0040-1707169.

(47) Gallucci, R. R.; Going, R. Chlorination of Aliphatic Ketones in Methanol. J. Org. Chem. 1981, 46 (12), 2532–2538. 10.1021/jo00325a019.

(48) Wang, J.; Li, H.; Zhang, D.; Huang, P.; Wang, Z.; Zhang, R.; Liang, Y.; Dong, D. Divergent Synthesis of α,α-Dihaloamides through α,α-Dihalogenation of β-Oxo Amides by Using N-Halosuccinimides. European Journal of Organic Chemistry 2013, 2013 (24), 5376–5380. 10.1002/ejoc.201300341.

(49) Tarantino, G.; Hammond, C. Catalytic C(Sp3)–F Bond Formation: Recent Achievements and Pertaining Challenges. Green Chem. 2020, 22 (16), 5195–5209. 10.1039/D0GC02067B.

(50) Ren, J.; Tong, R. Convenient in Situ Generation of Various Dichlorinating Agents from Oxone and Chloride: Diastereoselective Dichlorination of Allylic and Homoallylic Alcohol Derivatives. Org. Biomol. Chem. 2013, 11 (26), 4312–4315. 10.1039/C3OB40670A.

(51) Jin, J.; Zhao, Y.; Kyne, S. H.; Farshadfar, K.; Ariafard, A.; Chan, P. W. H. Copper(I)-Catalysed Site-Selective C(Sp3)–H Bond Chlorination of Ketones, (E)-Enones and Alkylbenzenes by Dichloramine-T. Nat Commun 2021, 12 (1), 4065. 10.1038/s41467-021-23988-y.

(52) Latham, J.; Henry, J.-M.; Sharif, H. H.; Menon, B. R. K.; Shepherd, S. A.; Greaney, M. F.; Micklefield, J. Integrated Catalysis Opens New Arylation Pathways via Regiodivergent Enzymatic C–H Activation. Nat Commun 2016, 7 (1), 11873. 10.1038/ncomms11873.

(53) Petrone, D. A.; Ye, J.; Lautens, M. Modern Transition-Metal-Catalyzed Carbon–Halogen Bond Formation. Chem. Rev. 2016, 116 (14), 8003–8104. 10.1021/acs.chemrev.6b00089.

(54) Xiong, H.-Y.; Cahard, D.; Pannecoucke, X.; Besset, T. Pd-Catalyzed Directed Chlorination of Unactivated C(Sp3)–H Bonds at Room Temperature. European Journal of Organic Chemistry 2016, 2016 (21), 3625–3630. 10.1002/ejoc.201600600.

(55) Li, G.; Dilger, A. K.; Cheng, P. T.; Ewing, W. R.; Groves, J. T. Selective C−H Halogenation with a Highly Fluorinated Manganese Porphyrin. Angewandte Chemie International Edition 2018, 57 (5), 1251–1255. 10.1002/anie.201710676.

(56) Ozawa, J.; Kanai, M. Silver-Catalyzed C(Sp3)–H Chlorination. Org. Lett. 2017, 19 (6), 1430–1433. 10.1021/acs.orglett.7b00367.

(57) Fawcett, A.; Keller, M. J.; Herrera, Z.; Hartwig, J. F. Site Selective Chlorination of C(Sp3)−H Bonds Suitable for Late-Stage Functionalization. Angewandte Chemie International Edition 2021, 60 (15), 8276–8283. 10.1002/anie.202016548.

(58) Quinn, R. K.; Könst, Z. A.; Michalak, S. E.; Schmidt, Y.; Szklarski, A. R.; Flores, A. R.; Nam, S.; Horne, D. A.; Vanderwal, C. D.; Alexanian, E. J. Site-Selective Aliphatic C–H Chlorination Using N-Chloroamides Enables a Synthesis of Chlorolissoclimide. J. Am. Chem. Soc. 2016, 138 (2), 696–702. 10.1021/jacs.5b12308.

(59) Smithwick, E. R.; Wilson, R. H.; Chatterjee, S.; Pu, Y.; Dalluge, J. J.; Damodaran, A. R.; Bhagi-Damodaran, A. Electrostatically Regulated Active Site Assembly Governs Reactivity in Nonheme Iron Halogenases. ACS Catal. 2023, 13 (20), 13743–13755. 10.1021/acscatal.3c02531.

(60) Martinie, R. J.; Livada, J.; Chang, W.; Green, M. T.; Krebs, C.; Bollinger, J. M.; Silakov, A. Experimental Correlation of Substrate Position with Reaction Outcome in the Aliphatic Halogenase, SyrB2. J. Am. Chem. Soc. 2015, 137 (21), 6912–6919. 10.1021/jacs.5b03370.

(61) Chen, Y.-H.; Comeaux, L. M.; Herbst, R. W.; Saban, E.; Kennedy, D. C.; Maroney, M. J.; Knapp, M. J. Coordination Changes and Auto-Hydroxylation of FIH-1: Uncoupled O2-Activation in a Human Hypoxia Sensor. Journal of Inorganic Biochemistry 2008, 102 (12), 2120–2129. 10.1016/j.jinorgbio.2008.07.018.

(62) Ryle, M. J.; Koehntop, K. D.; Liu, A.; Que, L.; Hausinger, R. P. Interconversion of Two Oxidized Forms of Taurine/α-Ketoglutarate Dioxygenase, a Non-Heme Iron Hydroxylase: Evidence for Bicarbonate Binding. PNAS 2003, 100 (7), 3790–3795. 10.1073/pnas.0636740100.

(63) Wong, S. D.; Srnec, M.; Matthews, M. L.; Liu, L. V.; Kwak, Y.; Park, K.; Bell III, C. B.; Alp, E. E.; Zhao, J.; Yoda, Y.; Kitao, S.; Seto, M.; Krebs, C.; Bollinger, J. M.; Solomon, E. Elucidation of the Fe(Iv)=O Intermediate in the Catalytic Cycle of the Halogenase SyrB2. Nature 2013, 499 (7458), 320–323. 10.1038/nature12304.

(64) Neugebauer, M. E.; Kissman, E. N.; Marchand, J. A.; Pelton, J. G.; Sambold, N. A.; Millar, D. C.; Chang, M. C. Y. Reaction Pathway Engineering Converts a Radical Hydroxylase into a Halogenase. Nat Chem Biol 2021, 1–9. 10.1038/s41589-021-00944-x.

(65) Liu, J.; Chakraborty, S.; Hosseinzadeh, P.; Yu, Y.; Tian, S.; Petrik, I.; Bhagi, A.; Lu, Y. Metalloproteins Containing Cytochrome, Iron–Sulfur, or Copper Redox Centers. Chem. Rev. 2014, 114 (8), 4366–4469. 10.1021/cr400479b.

(66) Mehmood, R.; Qi, H. W.; Steeves, A. H.; Kulik, H. J. The Protein’s Role in Substrate Positioning and Reactivity for Biosynthetic Enzyme Complexes: The Case of SyrB2/SyrB1. ACS Catal. 2019, 9 (6), 4930–4943. 10.1021/acscatal.9b00865.

(67) Kulik, H. J.; Drennan, C. L. Substrate Placement Influences Reactivity in Non-Heme Fe(II) Halogenases and Hydroxylases *. Journal of Biological Chemistry 2013, 288 (16), 11233–11241. 10.1074/jbc.M112.415570.

(68) Mehmood, R.; Vennelakanti, V.; Kulik, H. J. Spectroscopically Guided Simulations Reveal Distinct Strategies for Positioning Substrates to Achieve Selectivity in Nonheme Fe(II)/α-Ketoglutarate-Dependent Halogenases. ACS Catal. 2021, 11 (19), 12394–12408. 10.1021/acscatal.1c03169.

(69) Rugg, G.; M Senn, H. Formation and Structure of the Ferryl [Fe[Double Bond, Length as m-Dash]O] Intermediate in the Non-Haem Iron Halogenase SyrB2: Classical and QM/MM Modelling Agree. Physical Chemistry Chemical Physics 2017, 19 (44), 30107–30119. 10.1039/C7CP05937J.

(70) Huang, J.; Li, C.; Wang, B.; Sharon, D. A.; Wu, W.; Shaik, S. Selective Chlorination of Substrates by the Halogenase SyrB2 Is Controlled by the Protein According to a Combined Quantum Mechanics/Molecular Mechanics and Molecular Dynamics Study. ACS Catal. 2016, 6 (4), 2694–2704. 10.1021/acscatal.5b02825.

(71) Mitchell, A. J.; Zhu, Q.; Maggiolo, A. O.; Ananth, N. R.; Hillwig, M. L.; Liu, X.; Boal, A. Structural Basis for Halogenation by Iron- and 2-Oxo-Glutarate-Dependent Enzyme WelO5. Nat Chem Biol 2016, 12 (8), 636–640. 10.1038/nchembio.2112.

(72) Zhang, X.; Wang, Z.; Gao, J.; Liu, W. Chlorination versus Hydroxylation Selectivity Mediated by the Non-Heme Iron Halogenase WelO5. Physical Chemistry Chemical Physics 2020, 22 (16), 8699–8712. 10.1039/D0CP00791A.

(73) Nakamura, H.; Wang, J. X.; Balskus, E. P. Assembly Line Termination in Cylindrocyclophane Biosynthesis: Discovery of an Editing Type II Thioesterase Domain in a Type I Polyketide Synthase. Chem. Sci. 2015, 6 (7), 3816–3822. 10.1039/C4SC03132F.

(74) Zhang, W.; Ostash, B.; Walsh, C. T. Identification of the Biosynthetic Gene Cluster for the Pacidamycin Group of Peptidyl Nucleoside Antibiotics. Proceedings of the National Academy of Sciences 2010, 107 (39), 16828–16833. 10.1073/pnas.1011557107.

(75) Calderone, C. T.; Bumpus, S. B.; Kelleher, N. L.; Walsh, C. T.; Magarvey, N. A. A Ketoreductase Domain in the PksJ Protein of the Bacillaene Assembly Line Carries out Both α- and β-Ketone Reduction during Chain Growth. Proceedings of the National Academy of Sciences 2008, 105 (35), 12809–12814. 10.1073/pnas.0806305105.

(76) Vaillancourt, F. H.; Yeh, E.; Vosburg, D. A.; O’Connor, S. E.; Walsh, C. T. Cryptic Chlorination by a Non-Haem Iron Enzyme during Cyclopropyl Amino Acid Biosynthesis. Nature 2005, 436 (7054), 1191–1194. 10.1038/nature03797.

(77) Matthews, M. L.; Neumann, C. S.; Miles, L. A.; Grove, T. L.; Booker, S. J.; Krebs, C.; Walsh, C. T.; Bollinger, J. M. Substrate Positioning Controls the Partition between Halogenation and Hydroxylation in the Aliphatic Halogenase, SyrB2. PNAS 2009, 106 (42), 17723–17728. 10.1073/pnas.0909649106.

(78) Matthews, M. L.; Chang, W.; Layne, A. P.; Miles, L. A.; Krebs, C.; Bollinger, J. M. Direct Nitration and Azidation of Aliphatic Carbons by an Iron-Dependent Halogenase. Nat Chem Biol 2014, 10 (3), 209–215. 10.1038/nchembio.1438.

(79) Matthews, M. L.; Krest, C. M.; Barr, E. W.; Vaillancourt, F. H.; Walsh, C. T.; Green, M. T.; Krebs, C.; Bollinger, J. M. Substrate-Triggered Formation and Remarkable Stability of the C−H Bond-Cleaving Chloroferryl Intermediate in the Aliphatic Halogenase, SyrB2. Biochemistry 2009, 48 (20), 4331–4343. 10.1021/bi900109z.

(80) Tian, J.; Liu, J.; Knapp, M.; Donnan, P. H.; Boggs, D. G.; Bridwell-Rabb, J. Custom Tuning of Rieske Oxygenase Reactivity. Nat Commun 2023, 14 (1), 5858. 10.1038/s41467-023-41428-x.

(81) Ghirlanda, G. Design of Membrane Proteins: Toward Functional Systems. Current Opinion in Chemical Biology 2009, 13 (5), 643–651. 10.1016/j.cbpa.2009.09.017.

(82) Tian, S.; Jones, S. M.; Jose, A.; Solomon, E. I. Chloride Control of the Mechanism of Human Serum Ceruloplasmin (Cp) Catalysis. J. Am. Chem. Soc. 2019, 141 (27), 10736–10743. 10.1021/jacs.9b03661.

(83) Joseph, C. A.; Maroney, M. J. Cysteine Dioxygenase: Structure and Mechanism. Chem. Commun. 2007, No. 32, 3338–3349. 10.1039/B702158E.

