## Supplementary material for "Controllable multi-halogenation of a non-native substrate by SyrB2 iron halogenase": Supp Info_multihalogenation

### Materials and Methods

#### Protein Expression

SyrB1, SyrB2, Sfp, and TycF were expressed using the same protocol as previously described.[1] BL21 (DE3) cells (Thermo Scientific) were transformed with the pET-28a(+) or pET-30(+) vector for protein expression. A primary culture (~50 mL, 2XYT broth, 0.05 mg/mL kanamycin) was inoculated with a colony containing the appropriate protein and grown at 37°C/250 RPM overnight. Secondary cultures (1 L, 2XYT broth, 0.05 mg/mL kanamycin) in 2.8L flasks were inoculated with 10 mL of the primary culture and grown at 37°C/220 RPM till an $OD_{600}$ = 0.6 – 0.8 was reached. A single drop of Antifoam 204 (Sigma) was added to each culture for oxygen diffusion. Cultures were cooled on ice baths for 15-20 minutes prior to induction with Isopropyl β-D-1-thiogalactopyranoside (IPTG, GoldBio) at a final concentration of 0.2 mM. Cells were then grown for 18 hr at 18°C/220 RPM, harvested by centrifugation, flash frozen in liquid nitrogen, and stored at -20°C till purification. The above protocol describes the expression for SyrB1, SyrB2, and Sfp proteins. For TycF, the protocol is identical except the expression time is 72 hr and the [IPTG] = 1 mM.

#### Protein Purification

MilliQ water from a Barnstead GenPure water filtration system (Thermo Scientific) with a resistivity of at least 18.2 MΩ·cm was used to prepare all buffers described herein. For all proteins, thawed cell pellets were resuspended with 5 mL/g cells of wash buffer [50 mM HEPES (free acid, DOT Scientific), 300 mM NaCl (Sigma: BioUltra), and 5 mM imidazole (Sigma: ReagentPlus), pH = 7.5] supplemented with 1 mM phenylmethylsulfonyl fluoride (Sigma, added from 100 mM stock solution in IPA). Cells were lysed by sonication and centrifuged (4°C, 20,000 RPM, 20 min) to pellet cell debris. The supernatant was filtered through 0.22 μm syringe filters and loaded onto a wash-buffer equilibrated 5 mL HisTrapFF column (Cytiva, 2 mL/min binding rate) using an AKTA Start protein purification system (Cytiva). After application, the column was washed with 15 CV of wash buffer to elute non-specific binding proteins. A gradient elution from 0% to 100% elution buffer (50 mM HEPES, 100 mM NaCl, 250 mM imidazole, pH = 7.5) was applied to elute the desired protein. For TycF, an isocratic elution was used by washing the column with 10 CV of 30% elution buffer then 10 CV of 100% elution buffer. Relevant protein fractions, as determined by SDS–PAGE analysis, were pooled. Sfp and TycF were dialyzed against 3 L of RXN buffer (20 mM HEPES, pH = 7.5) at 4°C overnight, filtered, concentrated using 10 kDa MWCO centrifugal filter units (Corning), flash frozen in liquid nitrogen, and stored at −80°C.

SyrB2 was dialyzed against 3 L of RXN buffer supplemented with 1 mM EDTA overnight to remove exogenous cations. Two more rounds of dialysis against RXN buffer were performed to remove EDTA. After filtration, SyrB2 was concentrated in 30 kDa MWCO centricons (Corning), and buffer exchanged against RXN buffer three more times to remove low-weight impurities. After aliquoting, the protein was flash frozen in liquid nitrogen and stored at -80°C. Details about the preparation of SyrB1 after IMAC chromatography are described in the next section. All protein concentrations were determined by the absorbance at 280 nm using molar extinction coefficients from the ProtParam tool.

**Phosphopantetheine (Ppt) and Amino Acid attachment to SyrB1**

After IMAC chromatography, SyrB1 was dialyzed against 3 L of RXN buffer supplemented with 1 mM EDTA overnight to remove exogenous cations. Two more rounds of dialysis against RXN buffer were performed to remove EDTA. After filtration, SyrB1 was concentrated in 30 kDa MWCO centricons. For attachment of the Ppt cofactor, SyrB1 (100 µM) was incubated with coenzyme A (1 mM, Sigma: cofactor for acyl transfer), MgSO4 (5 mM, Sigma: BioReagent), and phosphopantetheinyl transferase Sfp (5 µM) in RXN buffer for 90 min at room temperature. The reaction was concentrated to 5 - 7 mL and loaded into the superloop of an AKTA Pure protein purification system (Cytiva) equipped with a HiLoad 26/600 Superdex 200 pg column (Cytiva). After equilibration with RXN buffer, an injection was taken and the protein was eluted at a flow rate of 0.8 mL/min. Usually 2-3 injections were performed depending on the amount of protein present (max injection 2 mL of 500 µM protein).

Relevant fractions, as determined by SDS-PAGE, were pooled and concentrated in 30 kDa MWCO centrifugal filter units. To append amino acid to the protein, SyrB1-Ppt (100 µM) was incubated with MgSO4 (5 mM), ATP (10 mM, Sigma: Grade I, ≥99%, from microbial), and 10 mM of L-alpha-aminobutyric acid (L-Aba; acquired from Sigma except 3,3,$D_2$-L-Aba which was procured from C/D/N isotopes) in RXN buffer for 30-40 min at room temperature. After this step, SyrB1-Ppt-AA was cooled to 4°C and concentrated. After five rounds of buffer exchange against RXN buffer to remove excess components, the protein was aliquoted, flash frozen, and stored at -80°C.

**Halogenation assay for UPLC-MS studies**

Halogenation assays and AQC derivatization were performed as previously described.[1] SyrB2 (25-400 µM) was mixed into a RXN buffer solution containing 2OG (Sigma: 99%(T); 0-10 mM) and NaCl or NaBr (100 mM) (final concentrations given in parentheses; final volume = 100 µL; all solutions prepared from substrate stocks prepared in RXN buffer). For reactions employing the ferredoxin (Fdx)-based reduction system, NADPH (Abcam, 500 µM), Fdx (Sigma: lyophilized powder prepared in RXN buffer; 5 µM), and Fdx Reductase (Sigma: lyophilized powder prepared in RXN buffer; 3 µM) were included. Then, ferrous ammonium sulfate (Sigma: BioUltra; 100 µM; added from a fresh stock in 2.5 mM $H_2SO_4$) was added followed immediately by SyrB1-Ppt-Aba (100 µM). Solutions were promptly mixed after each substrate addition and then incubated on a Thermomixer (Eppendorf) at 25°C at 300 RPM for 30 min. After incubation, the reactions were concentrated in 0.5 mL centrifugal filter units (Millipore-Sigma, 10 kDa MWCO). Reactions were buffer exchanged against RXN buffer five times to remove excess salt and reagents. After dilution to 125 µL, TycF thioesterase was added to a final concentration of 5 µM. The thioesterase reaction was allowed to proceed for 90 min. at 25°C before dilution to 400 µL with RXN buffer. The reactions were concentrated again and the flow-through was collected into deactivated, silanized vials (Waters). The dilution/concentration cycle was repeated 2x for flow-through collection.

Reaction flow-through was then lyophilized overnight and stored at -20°C till further use. Solid products were reconstituted in 178 µL of borate buffer (12.5 mM, pH = 8.5) and the pH was adjusted with 1.5 µL of 5 M NaOH. As an internal standard, 10 µL of 100 µM $^{13}C_4$-Thr (Cambridge Isotope Laboratories) was added to each solution. After reconstitution, 20 µL of 6-aminoquinolyl-N-hydroxysuccinimidyl carbamate (10 mM in acetonitrile, AQC, Cayman Chemical) was added to the solution, vortexed, and incubated at room temperature for 10 minutes. Samples were then analyzed by MS on the same day as derivatization or stored at -80°C till analysis.

Samples were analyzed using Sciex Exion UPLC coupled to a Sciex X500R quadrupole time-of-flight (qtof) for high-resolution MS or a Waters TQD UPLC/Triple Quadrupole-MS for MRM-MS. Separation methodologies were identical to those described previously.[1]

Percent conversions were calculated based on the MRM-MS data. Due to the unavailability of verified standards, we assumed that all species have the same ionization efficiency, thus producing identical signals for identical analyte concentrations. For any specific analyte, %conversion was calculated using the following formula:

$$\%\, product\ conversion = \frac{\sum specific\ analyte\ isotopic\ areas}{\sum all\ analytes'\, isotopic\ areas} * 100$$

For example, the %conversion for $Cl_2$ Aba would be equal to the integrated peak areas of $^{35}Cl^{35}Cl$-Aba + $^{35}Cl^{37}Cl$-Aba + $^{37}Cl^{37}Cl$-Aba divided by the integrated areas of unreacted Aba, the integrated area for lactonized Aba, the integrated areas of the isotopes of Cl-Aba ($^{35}Cl$-Aba + $^{37}Cl$-Aba), areas of the isotopes of $Cl_2$-Aba (given above), and the integrated areas of the isotopes of $Cl_3$-Aba ($^{35}Cl^{35}Cl^{35}Cl$-Aba + $^{35}Cl^{35}Cl^{37}Cl$-Aba + $^{35}Cl^{37}Cl^{37}Cl$-Aba + $^{37}Cl^{37}Cl^{37}Cl$-Aba). During all MRM runs, the masses for Aba, lactonized Aba, hydroxylated Aba, and mono-, di-, and tri-halogenated Aba were monitored. Chlorinated mass channels were monitored in chlorination reactions and brominated mass channels were monitored in bromination reactions. New product peaks were identified by comparison to MRM-MS control spectra with no added SyrB2 halogenase.

Mass error from q-ToF/MS was calculated using the following equation:

$$mass\ error\ (ppm) = \frac{(experimental\ mass - theoretical\ mass)}{theoretical\ mass} * 10^6$$

Simulated MS spectra for the compounds of interest were created using the Bruker DataAnalysis software.

### Molecular Dynamics simulations of SyrB2/SyrB1-Ppt-Aba

The starting structures were acquired from the SI of the work described by Kulik and coworkers.[2] To simulate the haloferryl intermediate, NO was removed and 2OG was truncated to succinate while the coordinating oxygen from the carboxylate functional group was treated as an oxo ligand. Force field parameters for the chloroferryl intermediate were created using the AMBER20 software's MCPB.py tool.[3,4] Geometry optimizations were performed with the Gaussian 16 software at the B3LYP/6-31G(d) level of theory while holding the alpha carbons of the protein ligands and the non-coordinating carboxylate carbon of succinate fixed.[5] Frequency calculations were performed with the identical level of theory. With the tleap module in AMBER20, SyrB1/2 proteins were described with the ff19SB force field.[6] The Ppt-Aba residue was parameterized according to the previous study.[2] Ppt-$Cl_2$-Aba was parameterized using a protocol as previously described.[7]

The solvent was treated as OPC water and solvated in a 10 Å box around the complex; counterions $Na^+$ and $Cl^-$ were added to neutralize the system. The complex was energy minimized (first solvent, then solvent and protein), gently heated to 300 K, and density equilibrated for 2 ns. Three independent 500 ns production runs were then executed. Since Kulik et al. previously showed that the AMBER force fields were inadequate for experimentally relevant conformational sampling, we applied identical substrate-based restraints during our simulations, with the exception that the N from the NO in their work was replaced with the oxo ligand in this study.[2] Trajectory analyses were performed with the CPPTRAJ module.[8] Error bars for the analyses were calculated using the standard error over the independent simulations (the standard deviation divided by $\sqrt{3}$).[9]

### Density Functional Theory Calculations

For geometry optimizations, truncated versions of Ppt-Aba were prepared in by capping the thioester with a methyl group. Geometry optimizations and were performed using the ORCA software (v. 5.0.3)[10] using the M06-2X/def2-TZVP level of theory[11,12]. The Resolution of Identity (RI) approximation was employed to accelerate coulombic integration and exchange chain of sphere exchange integrals (keywords: RIJCOSX; def2-J for calling auxiliary basis).[13] Calculations were performed in a dielectric of 4 to simulate the effect of the protein environment (CPCM solvation model[14]). Given the use of Minnesota Functionals, the integration grid sizes were increased from the default option (keyword: defgrid3). After optimization to minima (confirmed by frequency analysis showing zero imaginary frequencies), H atoms on $C_4$ were replaced by Cl atoms and the geometry optimizations and frequencies were repeated. Radicals were generated by deleting a single H-atom and reoptimizing as an open shell singlet. Bond Dissociation Energies were calculated from the change in electronic energies:

$$\mathrm{BDE\ (AA)} = \mathrm{E(\ H\text{-}atom + product\ radical)} - \mathrm{E(\ AA\ reactant)}$$

**Generation of Alkyne functional group from $Br_2$-Aba and Alkyne click assay**

A halogenation assay at a scale of 1 mL (100 µM SyrB1-Ppt-Aba/SyrB2/FAS, 10 mM 2OG, and 100 mM NaBr) was conducted as described above. Lyophilized dry stocks were reconstituted in 200 µL dioxane-water mixture containing xs Boc-anhydride/$Na_2CO_3$ for the protection of the amine group. After 1 hour of stirring at room temperature, 500 µL of a 1:1 water:chloroform mixture was added, and the solution was vortexed. The chloroform layer was collected and dried. A dipotassium salt of 1,3-propanediamine (final concentration: 250 µM) was prepared by stirring KH in 1,3-propanediamine for 1h at room temperature and cooled on ice prior to addition to amino acid sample. The amino acid sample was taken in 100 µL 1,3-propanediamine and to it ~5 µL dipotassium salt of 1,3-propanediamine added at 0°C. The mixture was then allowed to stir for 3h at room temperature. The reaction was diluted with 500 µL 1:1 water:chloroform and the organic layer was collected. After washing with 250 µL 1 M aq. HCl, the organic layer was collected and dried under $N_2$ gas. Dried product was reconstituted in 10 µL of methanol, vortexed, centrifuged, and taken for use in the click reaction assay described below.

A master mix for the click assay was prepared in the dark by mixing 25 µL of PBS (phosphate buffered saline, pH = 7.5), 25 µL 1 mM BTTAA (in MQ $H_2O$, Sigma), 25 µL 50 mM sodium ascorbate (dissolved in PBS pH = 7.5, Sigma: BioXtra), 25 µL 500 µM $CuSO_4 \cdot 5 H_2O$ (in MQ $H_2O$, Sigma: ACS Reagent), and 25 µL of 100 µM CalFluor 488 Azide (in DMSO, ClickChemistryTools). To a solid black 384-well plate (Griener), 10 µL of the click master mix plus 10 µL of standard L-Homopropargylglycine (HPG, ClickChemistryTools) or reaction-generated alkyne were added and mixed into individual wells. The plate was loaded onto a Tecan Spark plate reader instrument and shaken for 15 minutes at room temperature prior to recording the fluorescence ($\lambda_{ex}$ = 485 nm; $\lambda_{em}$ = 528 nm).

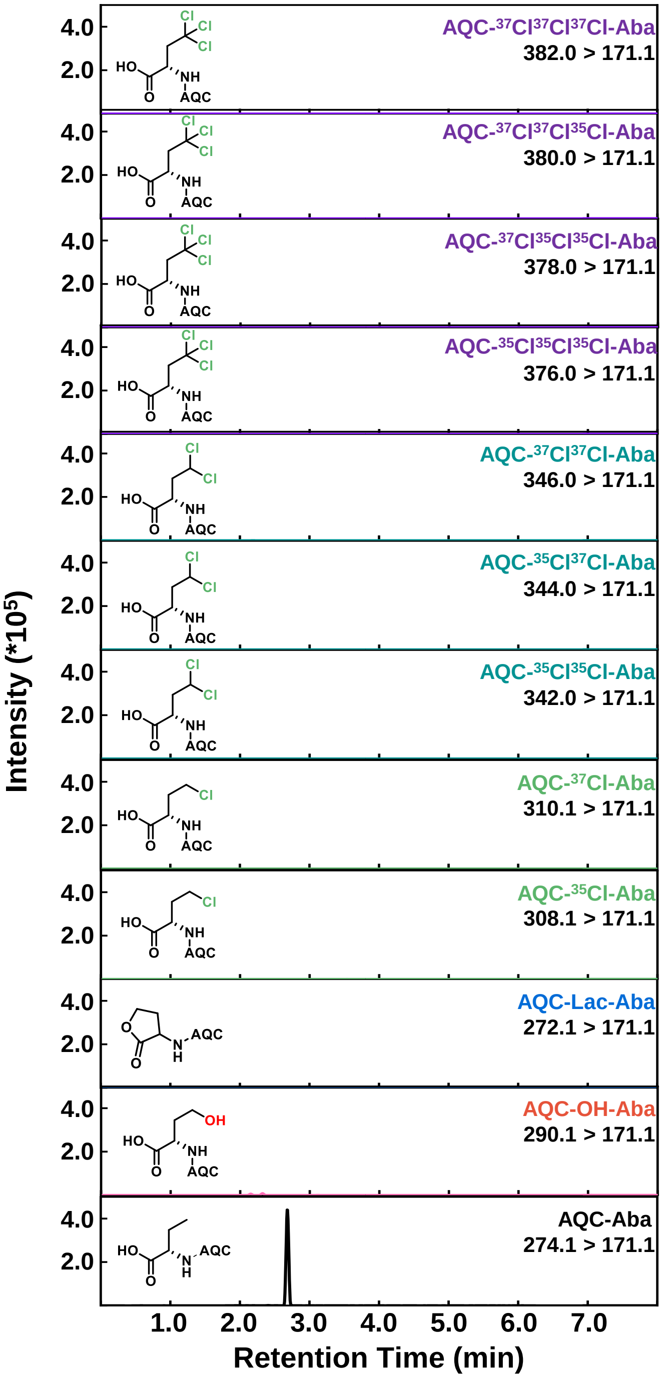


**Supplementary Figure 1.** Representative UPLC-MRM chromatogram for the control chlorination reaction of SyrB1-Ppt-Aba with no SyrB2 halogenase.

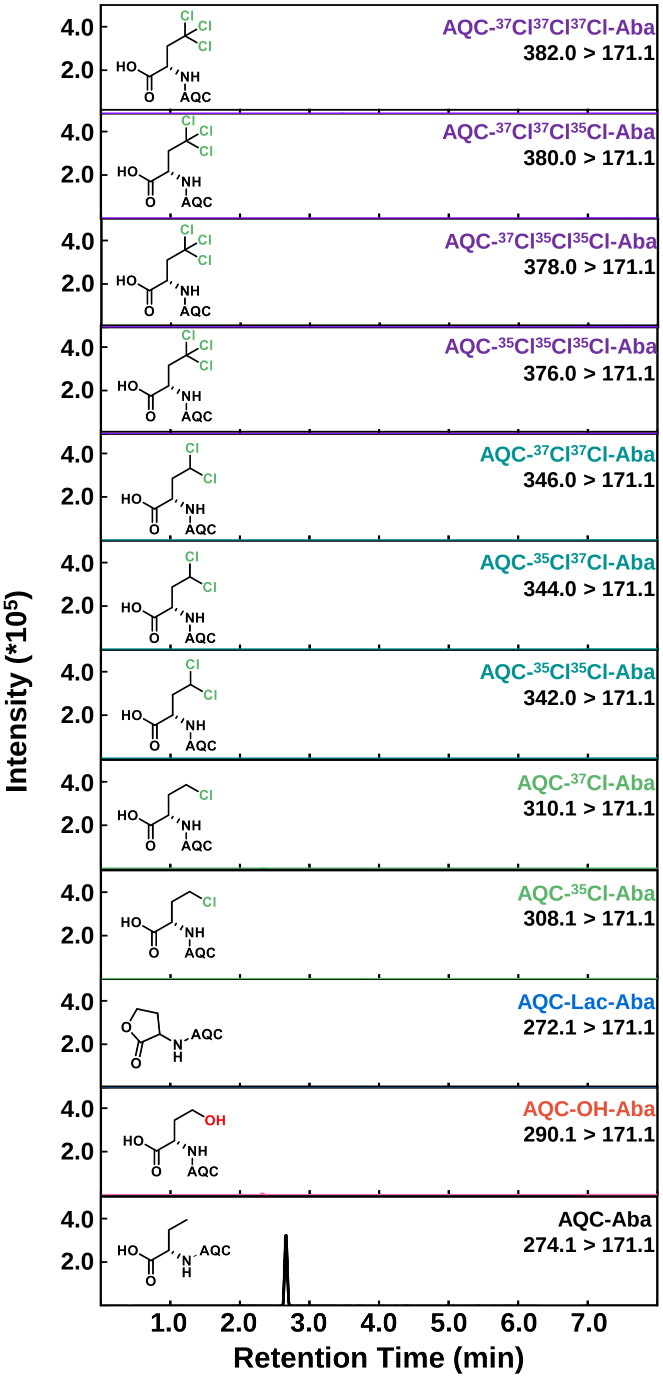


**Supplementary Figure 2.** Representative UPLC-MRM chromatogram for the chlorination reaction of SyrB1-Ppt-Aba with 0 mM 2OG.

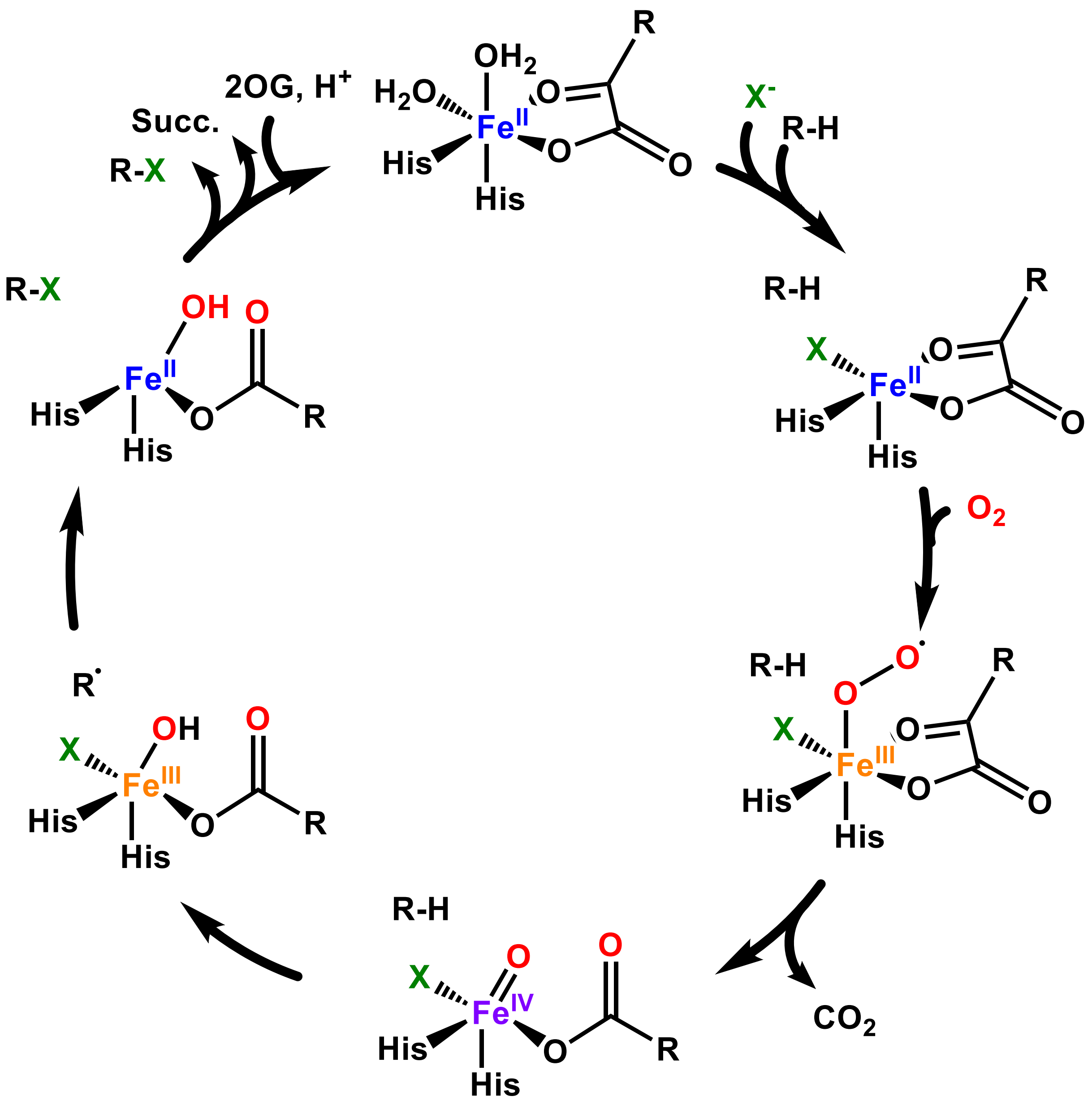


**Supplementary Figure 3.** Consensus mechanism for 2OG-dependent halogenases. After the halogenase binds ferrous iron and 2OG, the halide ligand binds to the iron center and substrate binds in the protein active site which triggers oxygen binding. After oxygen binds to the iron center, a decarboxylation of 2OG occurs which generates a high-valent haloferryl intermediate. This intermediate performs an H-atom abstraction on the substrate C-H bond forming a ferric hydroxyl species and a substrate radical. Halogenases position their substrates to optimize rebound with the halide ligand, which will generate a C-X bond (though rebound with the hydroxyl ligand is also possible, which forms a hydroxylated side product). After the halogenated product and succinate by-product leave the active site, the catalytic cycle can begin anew.

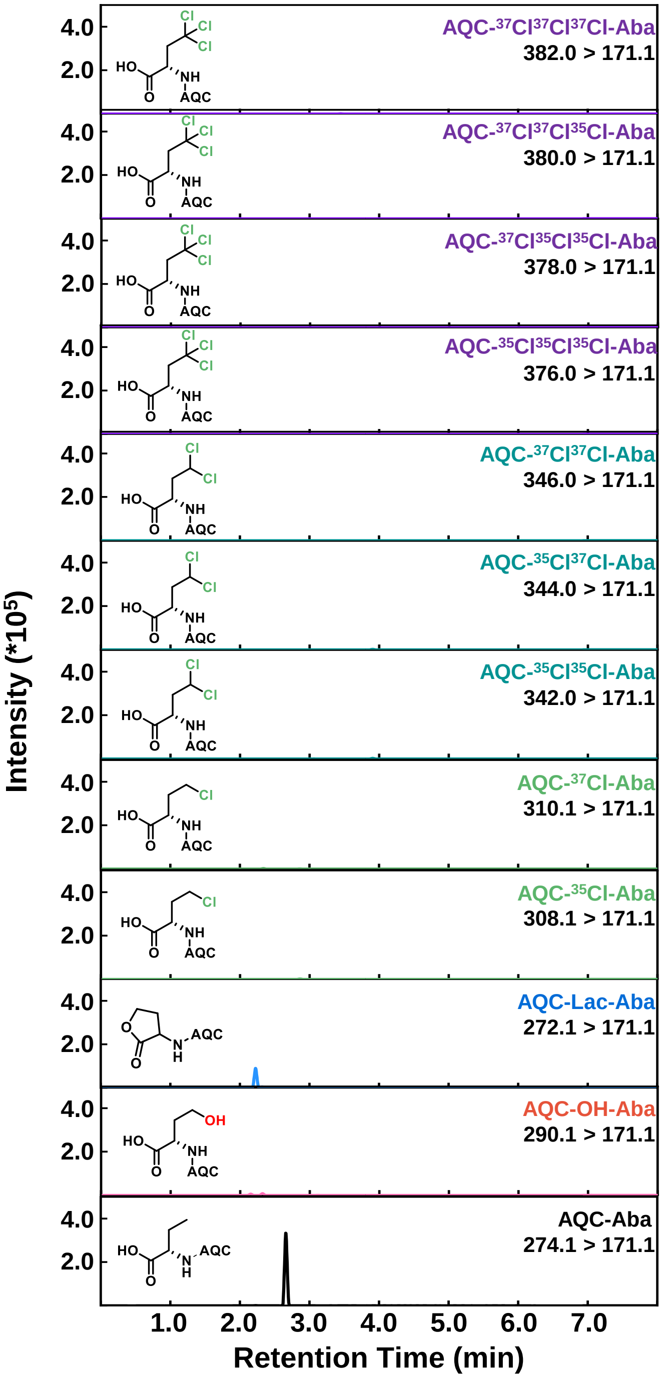


**Supplementary Figure 4.** Representative UPLC-MRM chromatogram for the chlorination reaction of SyrB1-Ppt-Aba with 50 µM 2OG.

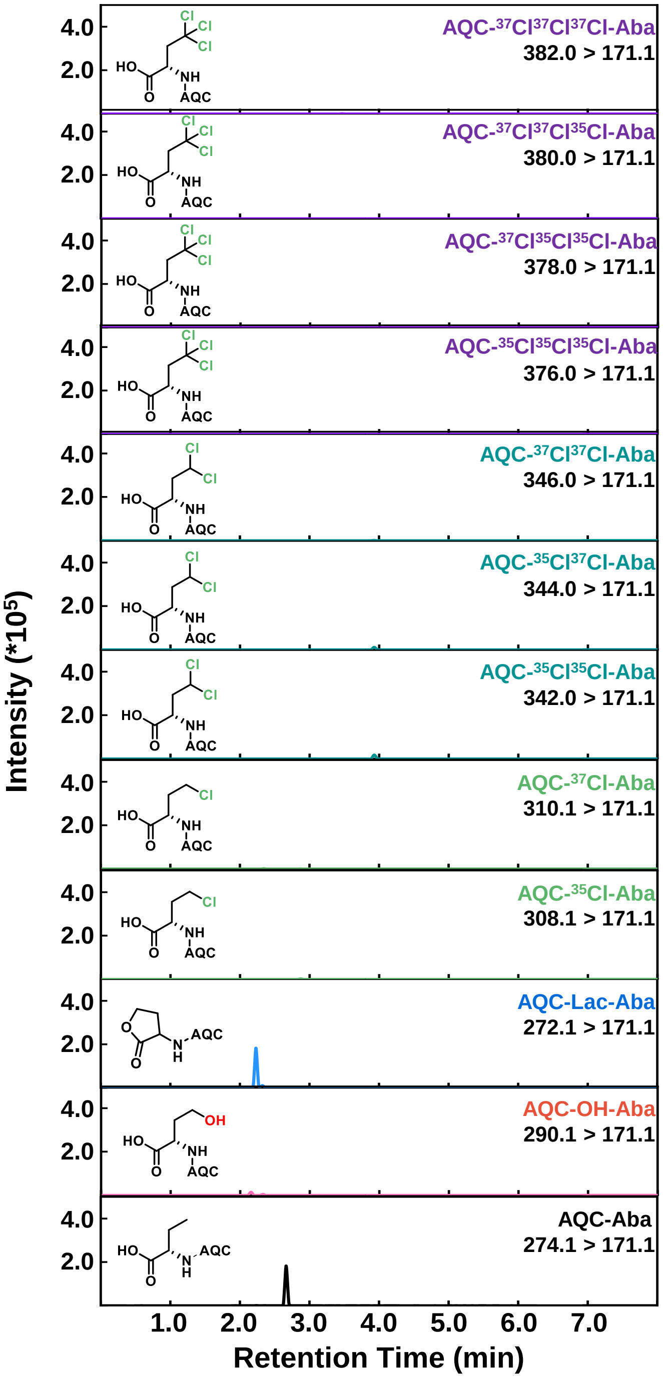


**Supplementary Figure 5.** Representative UPLC-MRM chromatogram for the chlorination reaction of SyrB1-Ppt-Aba with 100 µM 2OG.

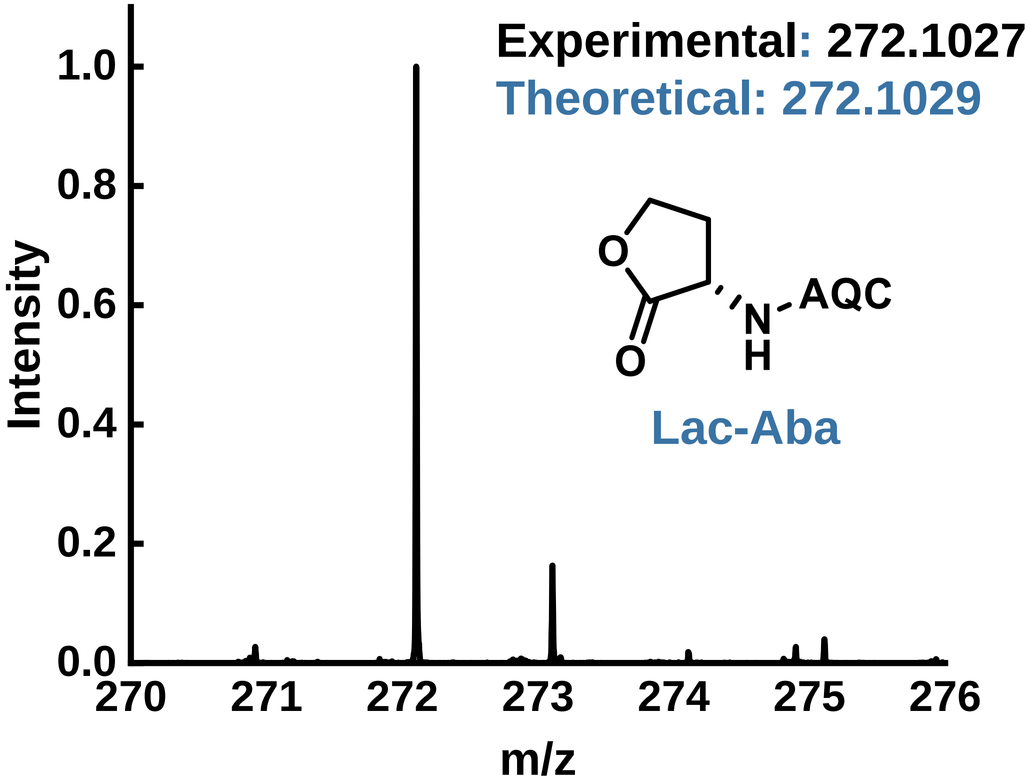


**Supplementary Figure 6.** Accurate mass spectrum validating the identity of lactonized Aba, generated by intramolecular $S_N2$ reaction. Mass Error = 1.1 ppm.

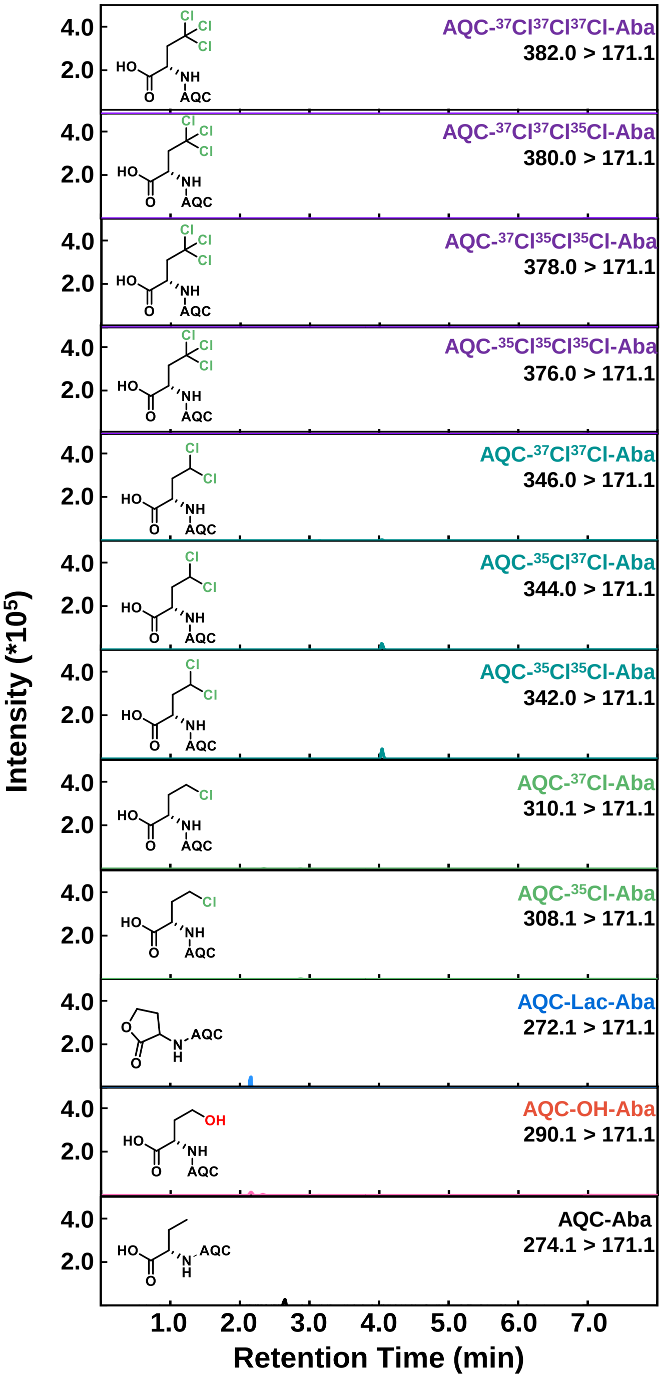


**Supplementary Figure 7.** Representative UPLC-MRM chromatogram for the chlorination reaction of SyrB1-Ppt-Aba with 250 µM 2OG.

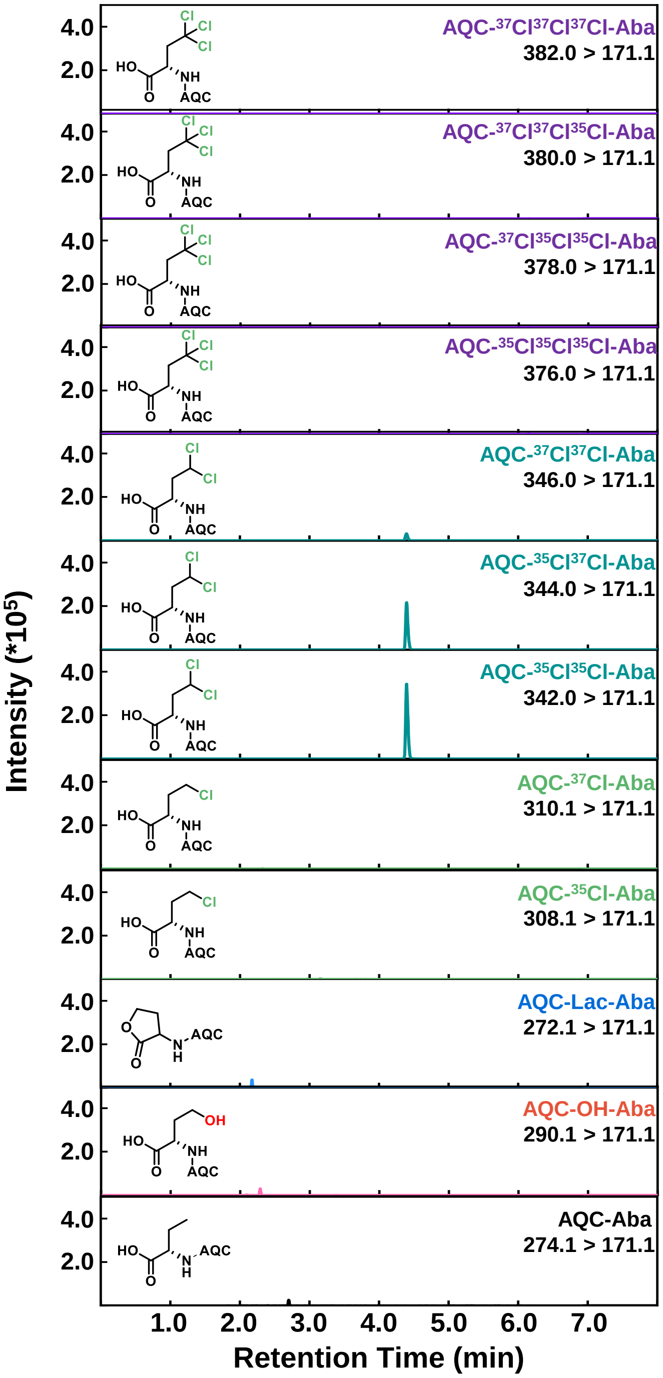


**Supplementary Figure 8.** Representative UPLC-MRM chromatogram for the chlorination reaction of SyrB1-Ppt-Aba with 500 μM 2OG.

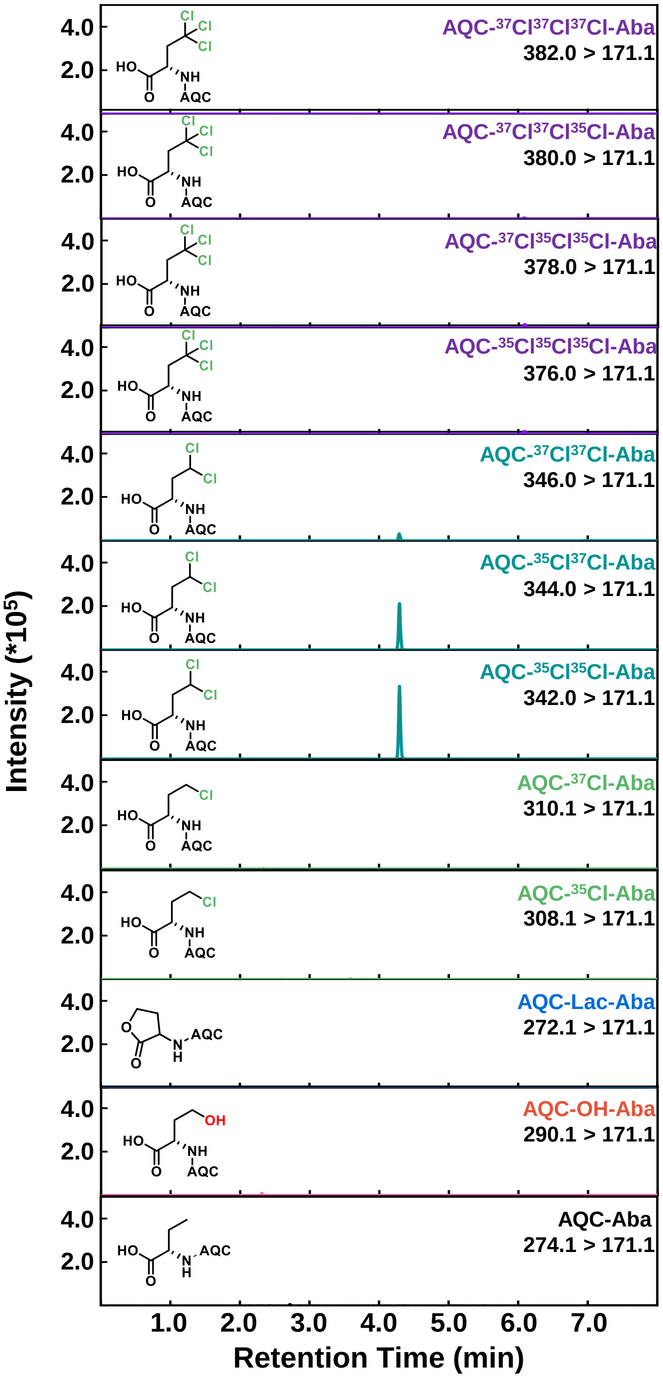


**Supplementary Figure 9.** Representative UPLC-MRM chromatogram for the chlorination reaction of SyrB1-Ppt-Aba with 1 mM 2OG.

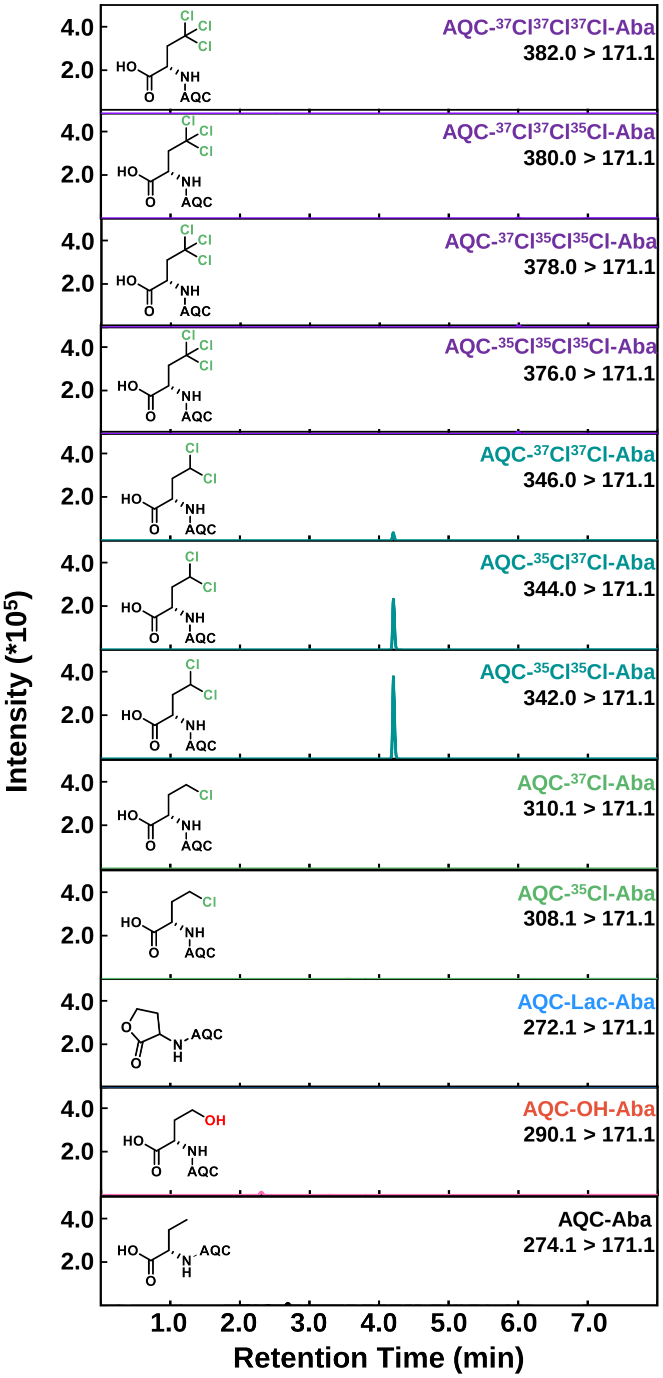


**Supplementary Figure 10.** Representative UPLC-MRM chromatogram for the chlorination reaction of SyrB1-Ppt-Aba with 10 mM 2OG.

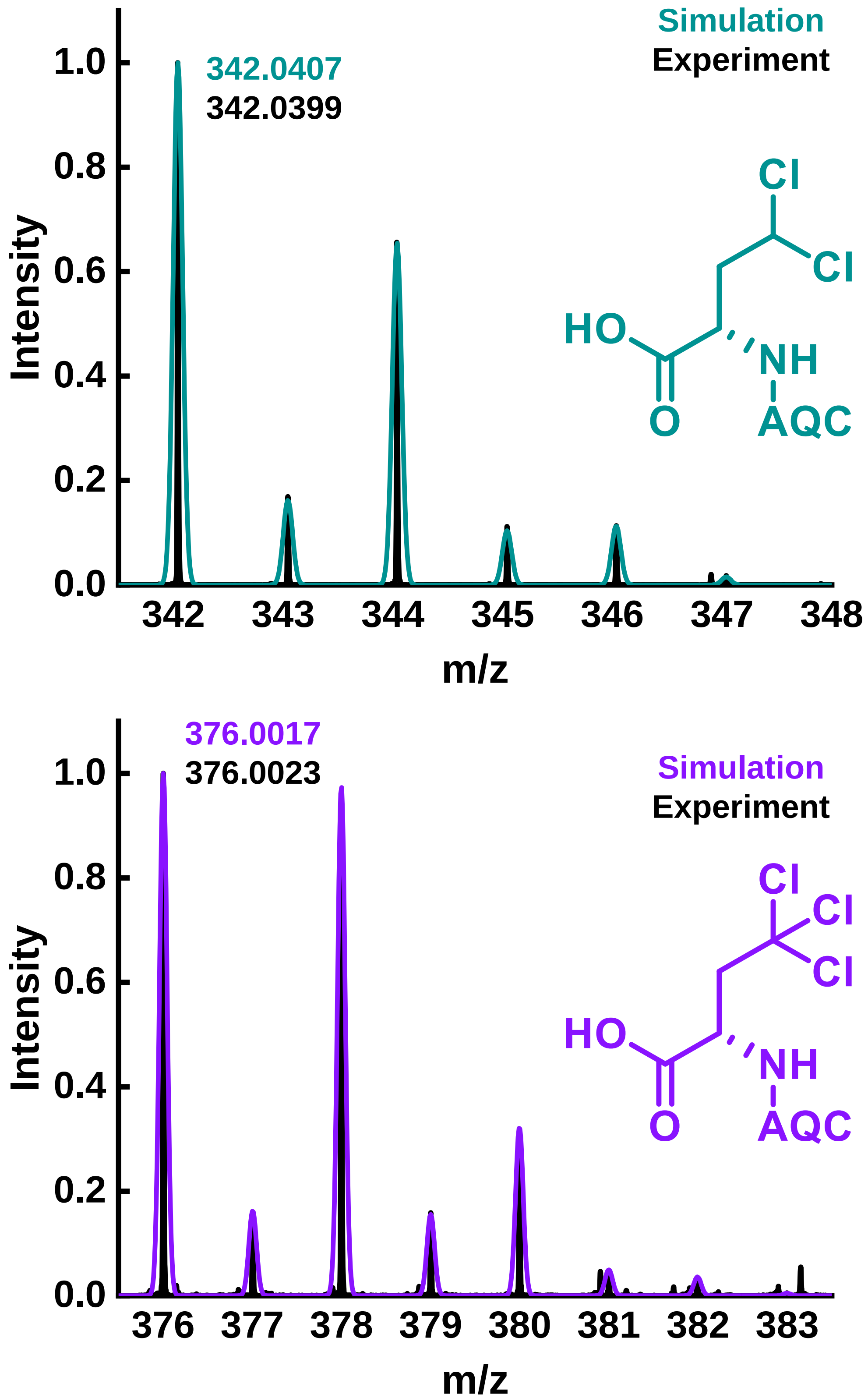


**Supplementary Figure 11.** Accurate mass spectra for analytes $Cl_2$-Aba (top) and $Cl_3$-Aba (bottom). Experimental masses strongly agree with the theoretical masses (error < 5 ppm), confirming analyte identity.

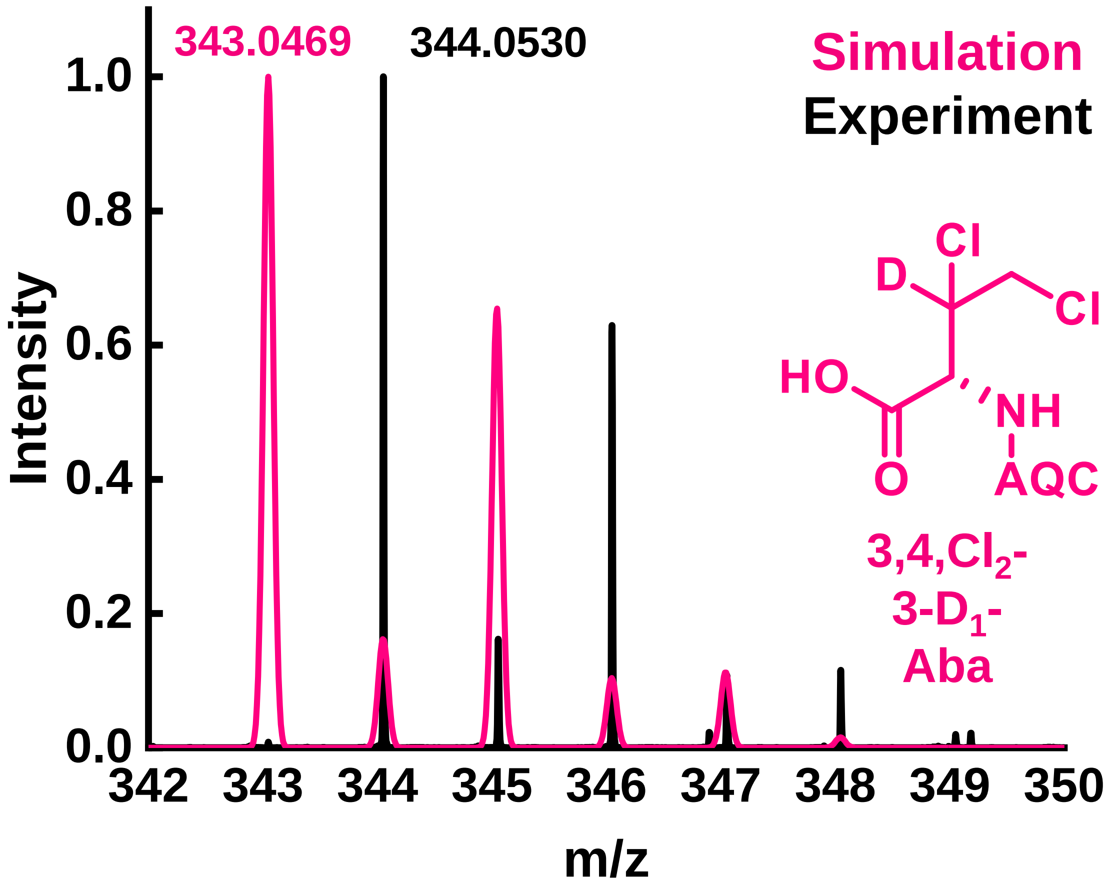


**Supplementary Figure 12.** Accurate mass spectrum for the di-chlorinated product from the chlorination reaction with SyrB1-Ppt-3,3-$D_2$-Aba overlayed with the simulated spectrum for 3,4,$Cl_2$-3-$D_1$-Aba.

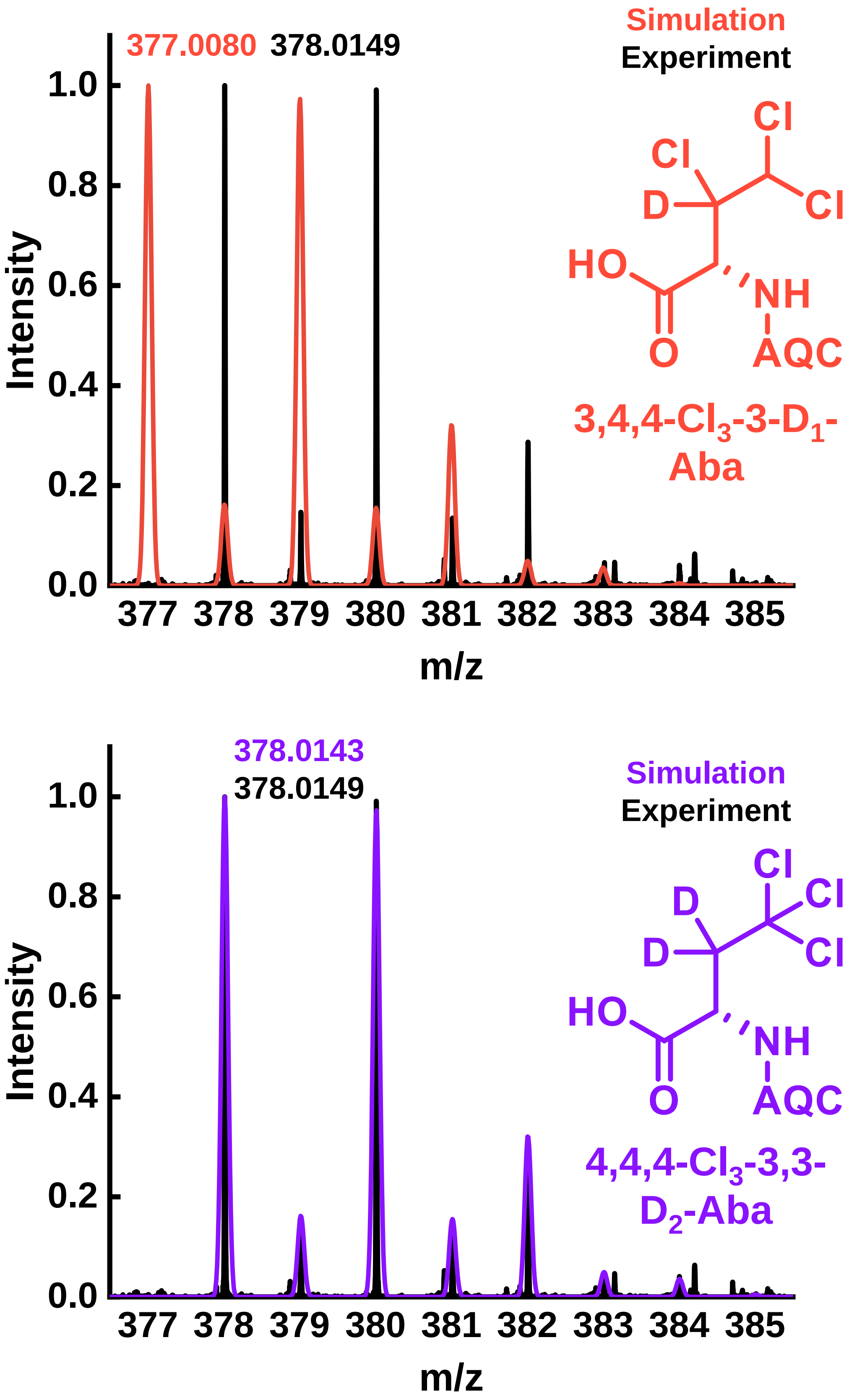


**Supplementary Figure 13.** Accurate mass spectra for the tr-chlorinated product with simulated 3,4,4-$Cl_3$-3-$D_1$-Aba (top) and 4,4,4-$Cl_3$-3,3-$D_2$-Aba (bottom). Isotope distribution pattern and experimental mass strongly agrees with the theoretical mass for 4,4,4-$Cl_3$-3,3-$d_2$-Aba (monoisotopic mass error = 1.6 ppm), confirming that both deuterium atoms were retained in the chlorination reaction, and all chlorine atoms were installed at C4.

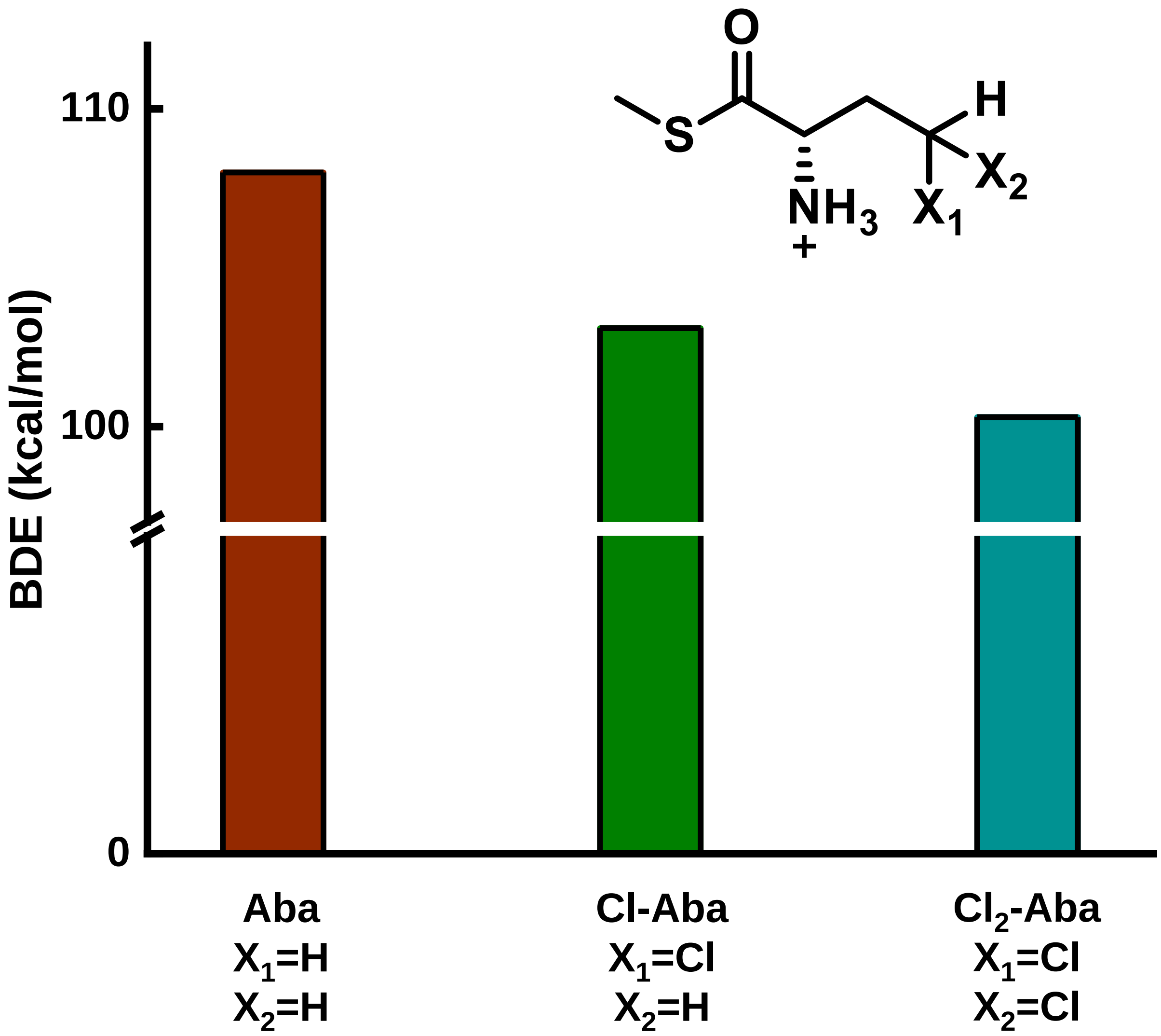


**Supplementary Figure 14.** DFT calculated bond dissociation energies (BDE, $C_4$-H bond) for H-abstraction at the $C_4$ position of Aba and chlorinated products shows that successive chlorination events reduce BDE, favoring multi-chlorination.

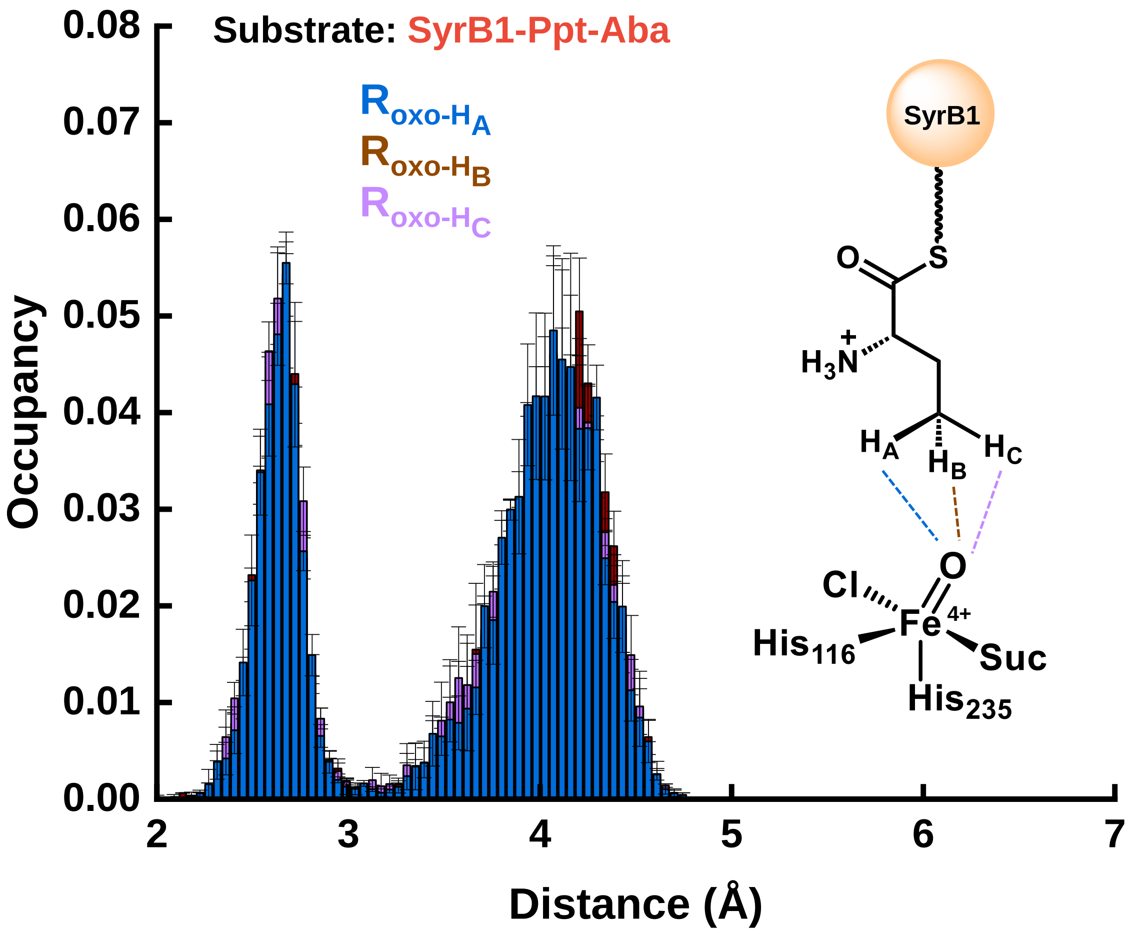


**Supplementary Figure 15.** MD simulation results monitoring the $C_4$ hydrogen atom distances of SyrB1-Ppt-Aba to the oxo ligand of the haloferryl intermediate in the SyrB2 active site. MD error bars are the standard error across three, independent production runs.

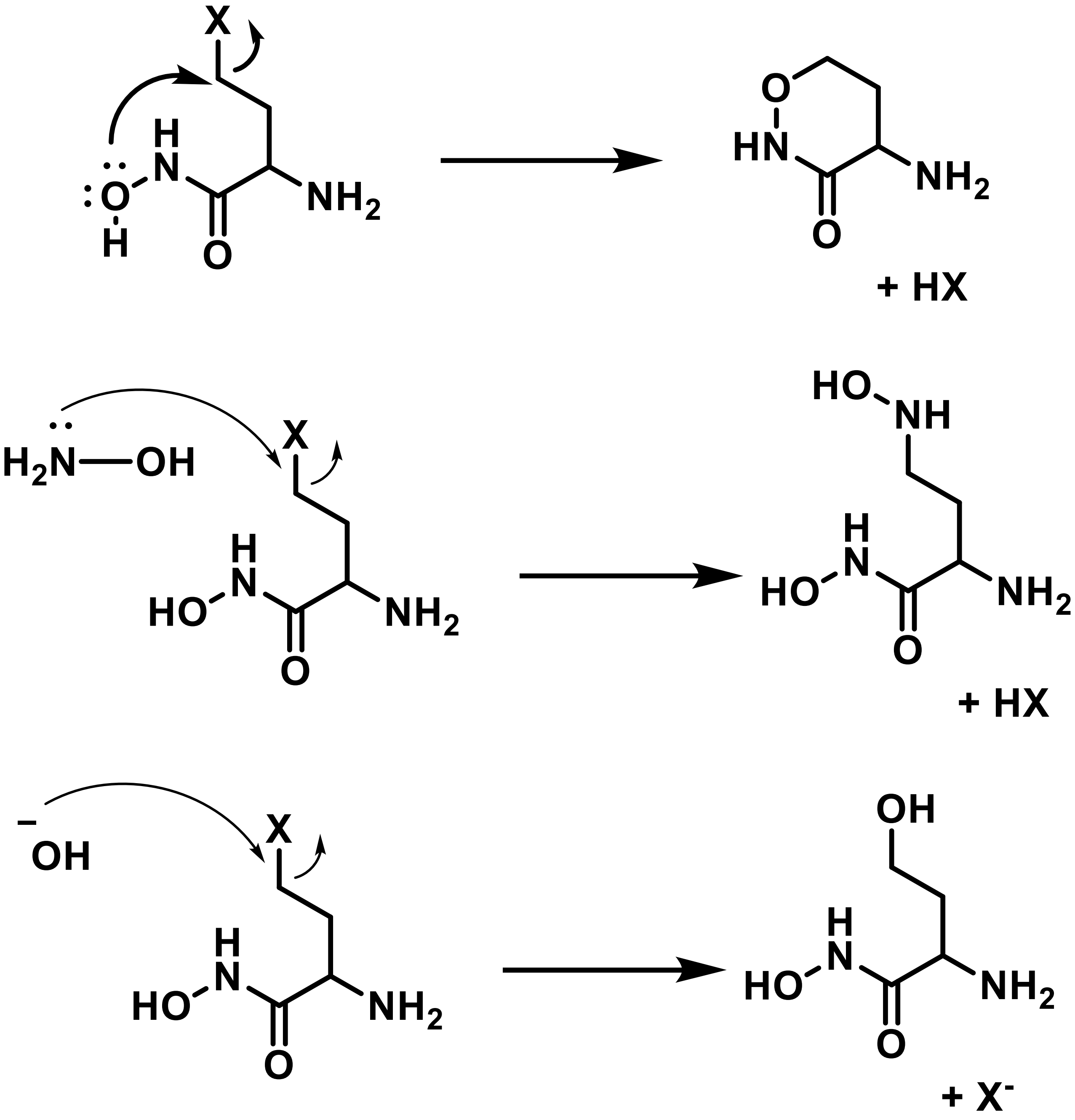


**Supplementary Figure 16.** Alternative reactions Cl-Aba can undergo under hydroxylamine derivatization conditions. Cyclization of halogenated hydroxymate derivatives of SyrB2 reaction products (top). Nucleophilic attack of hydroxamic acid analog of Cl-Aba by hydroxylamine (middle). Under basic conditions of hydroxylamine, mono-halogenated products can also be converted to hydroxylated products in a non-enzymatic fashion (bottom).

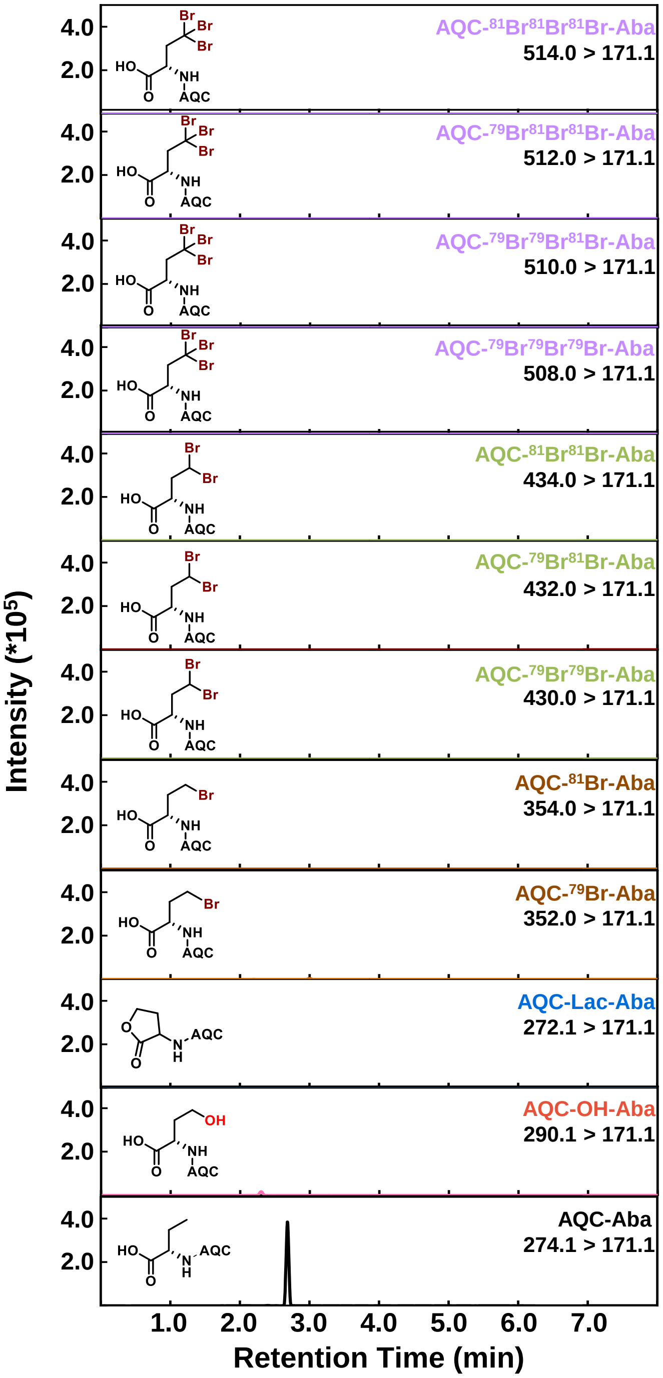


**Supplementary Figure 17.** Representative UPLC-MRM chromatogram for the control bromination reaction of SyrB1-Ppt-Aba with no SyrB2 halogenase.

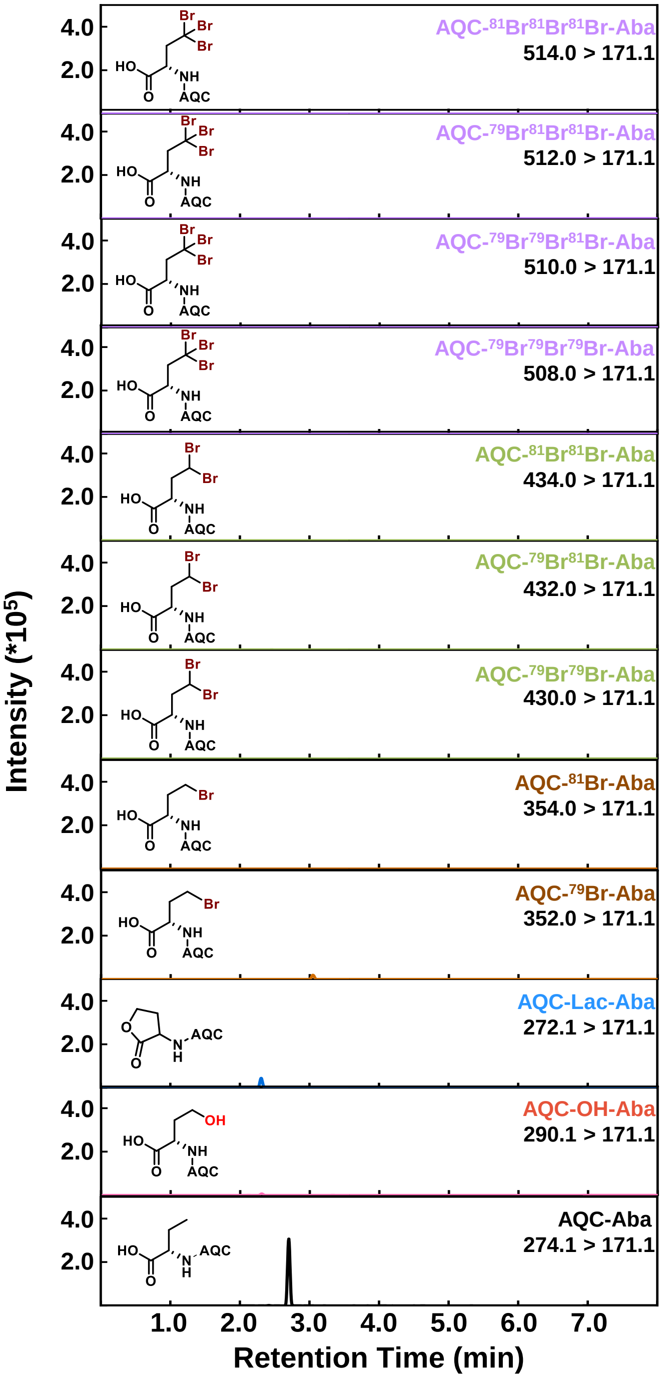


**Supplementary Figure 18.** Representative UPLC-MRM chromatogram for the bromination reaction of SyrB1-Ppt-Aba with 50 µM 2OG .

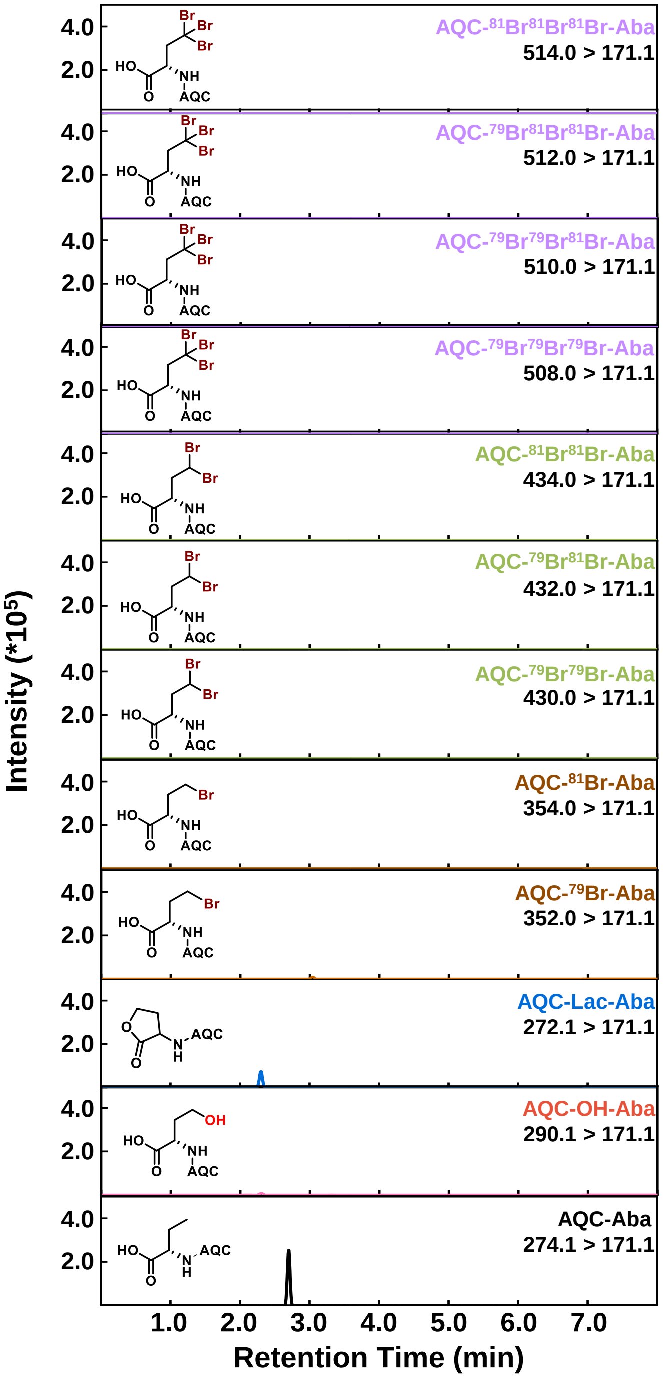


**Supplementary Figure 19.** Representative UPLC-MRM chromatogram for the bromination reaction of SyrB1-Ppt-Aba with 100 µM 2OG .

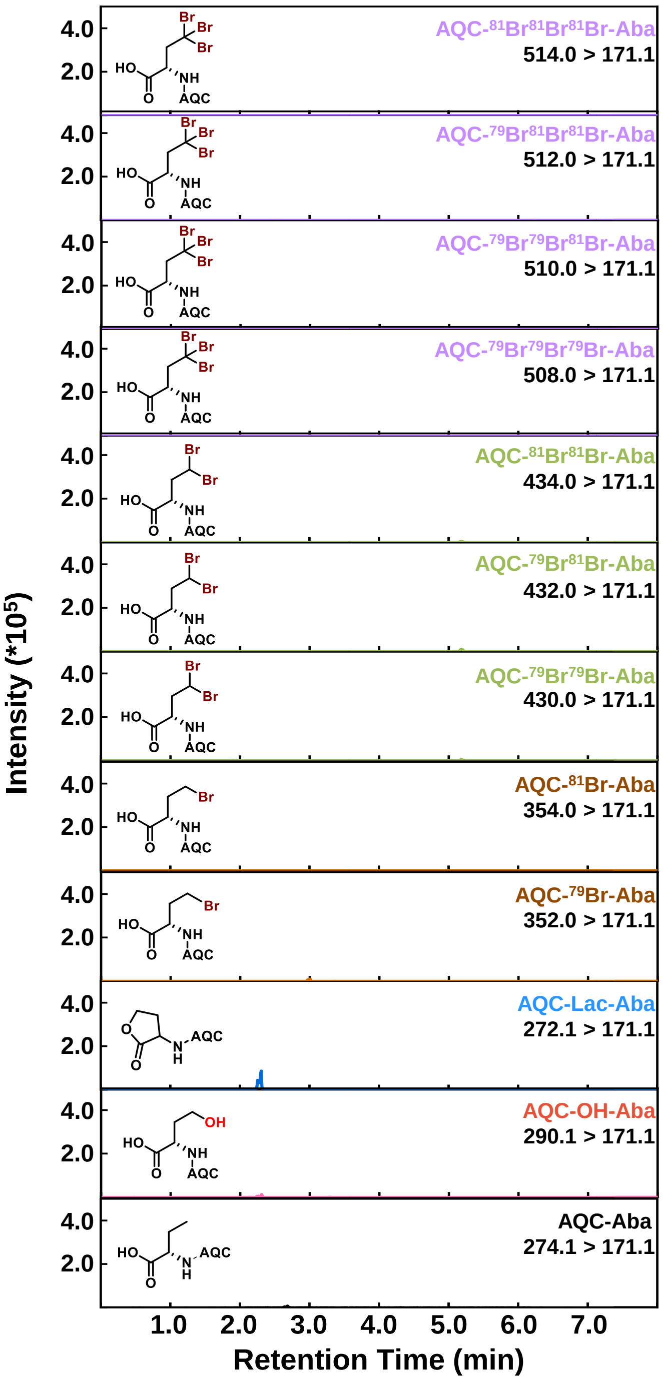


**Supplementary Figure 20.** Representative UPLC-MRM chromatogram for the bromination reaction of SyrB1-Ppt-Aba with 250 µM 2OG .

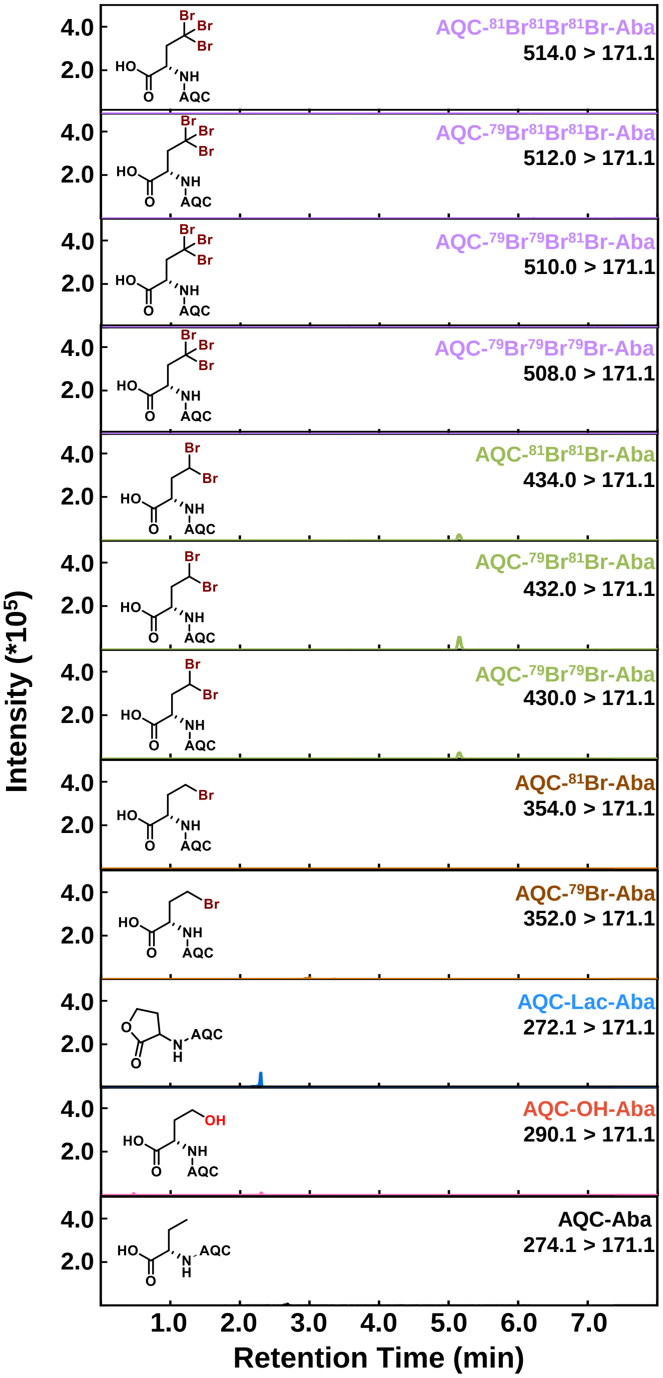


**Supplementary Figure 21.** Representative UPLC-MRM chromatogram for the bromination reaction of SyrB1-Ppt-Aba with 500 µM 2OG .

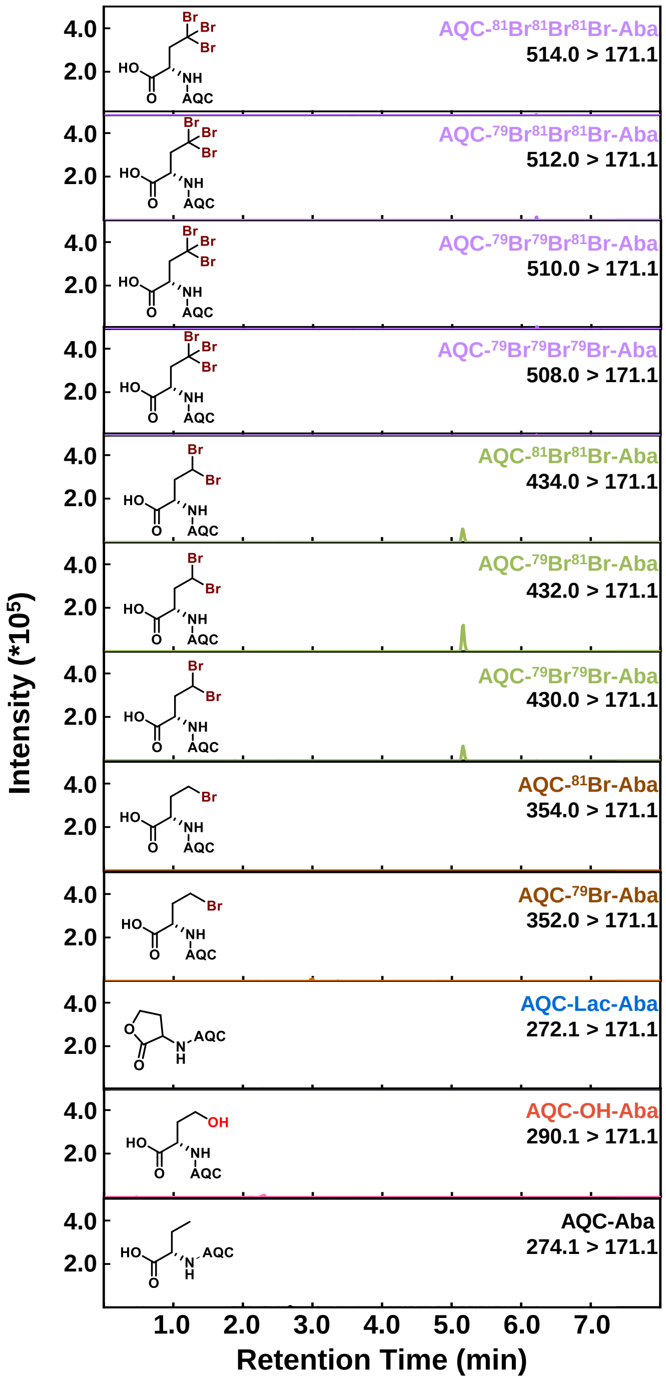


**Supplementary Figure 22.** Representative UPLC-MRM chromatogram for the bromination reaction of SyrB1-Ppt-Aba with 1 mM 2OG .

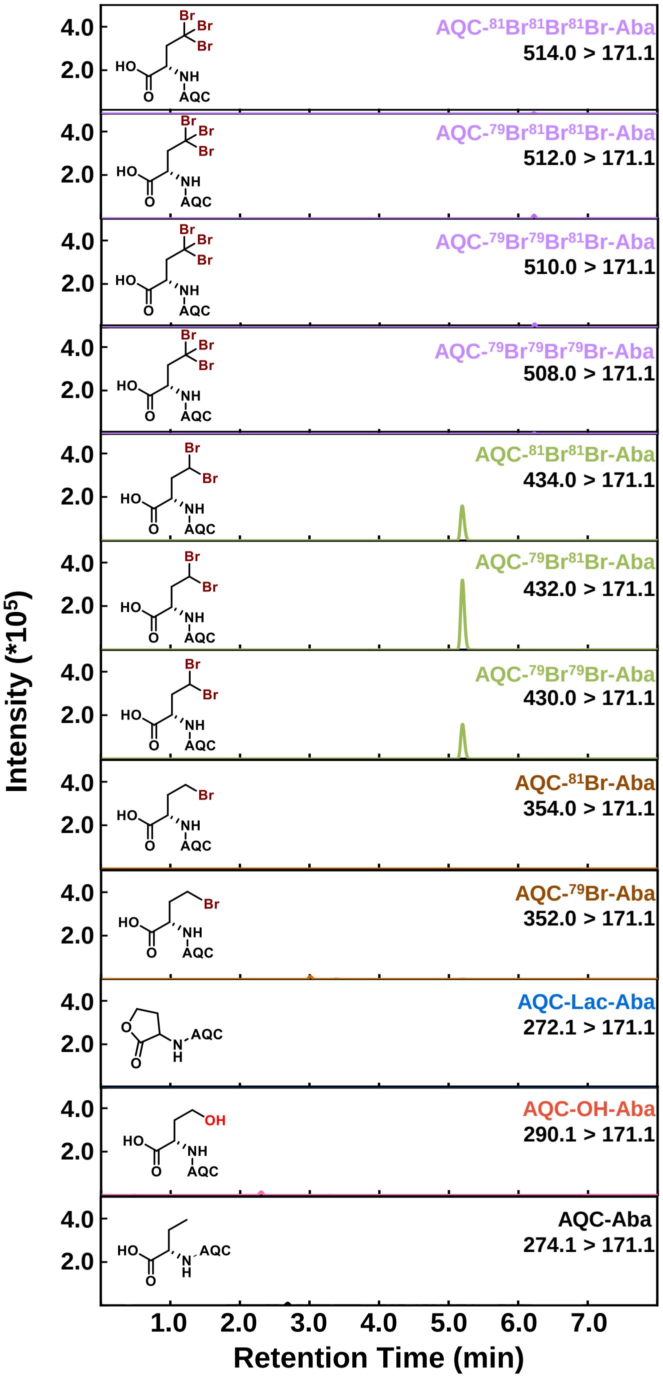


**Supplementary Figure 23.** Representative UPLC-MRM chromatogram for the bromination reaction of SyrB1-Ppt-Aba with 10 mM 2OG .

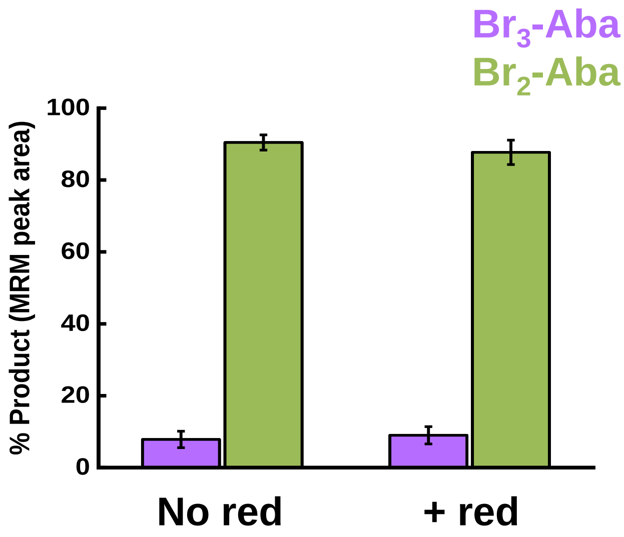


**Supplementary Figure 24.** AQC-tagged products detected as a percentage of total MRM peak area for SyrB2's reaction with SyrB1-Ppt-Aba with and without the biological reductant (abbreviated as red). The substrate to catalyst loading ratio was kept at 1:1 and the 2OG loading was set to 100 equivalents. Error bars are the standard deviation from n = 3 reactions. The % MRM peak area reflects trends in product selectivity and not definitive molar ratio.

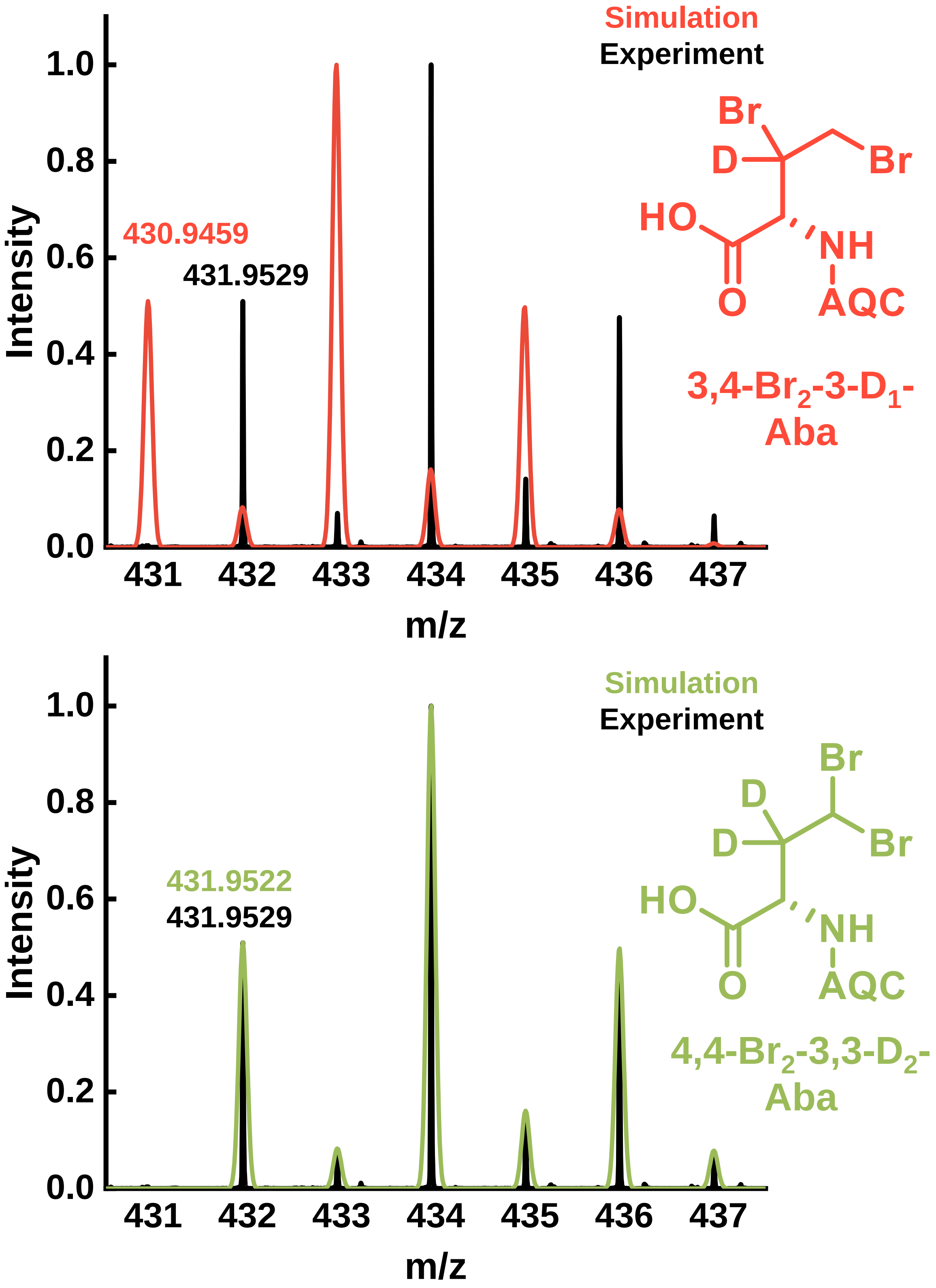


**Supplementary Figure 25.** Accurate mass spectra for the di-brominated product with simulated 3,4-$Br_2$-3-$D_1$-Aba (top) and 4,4-$Br_2$-3,3-$D_2$-Aba (bottom). Isotope distribution pattern and experimental mass strongly agrees with the theoretical mass for 4,4-$Br_2$-3,3-$D_2$-Aba (monoisotopic mass error = 1.6 ppm), confirming that both deuterium atoms were retained in the bromination reaction, and both bromine atoms were installed at C4.

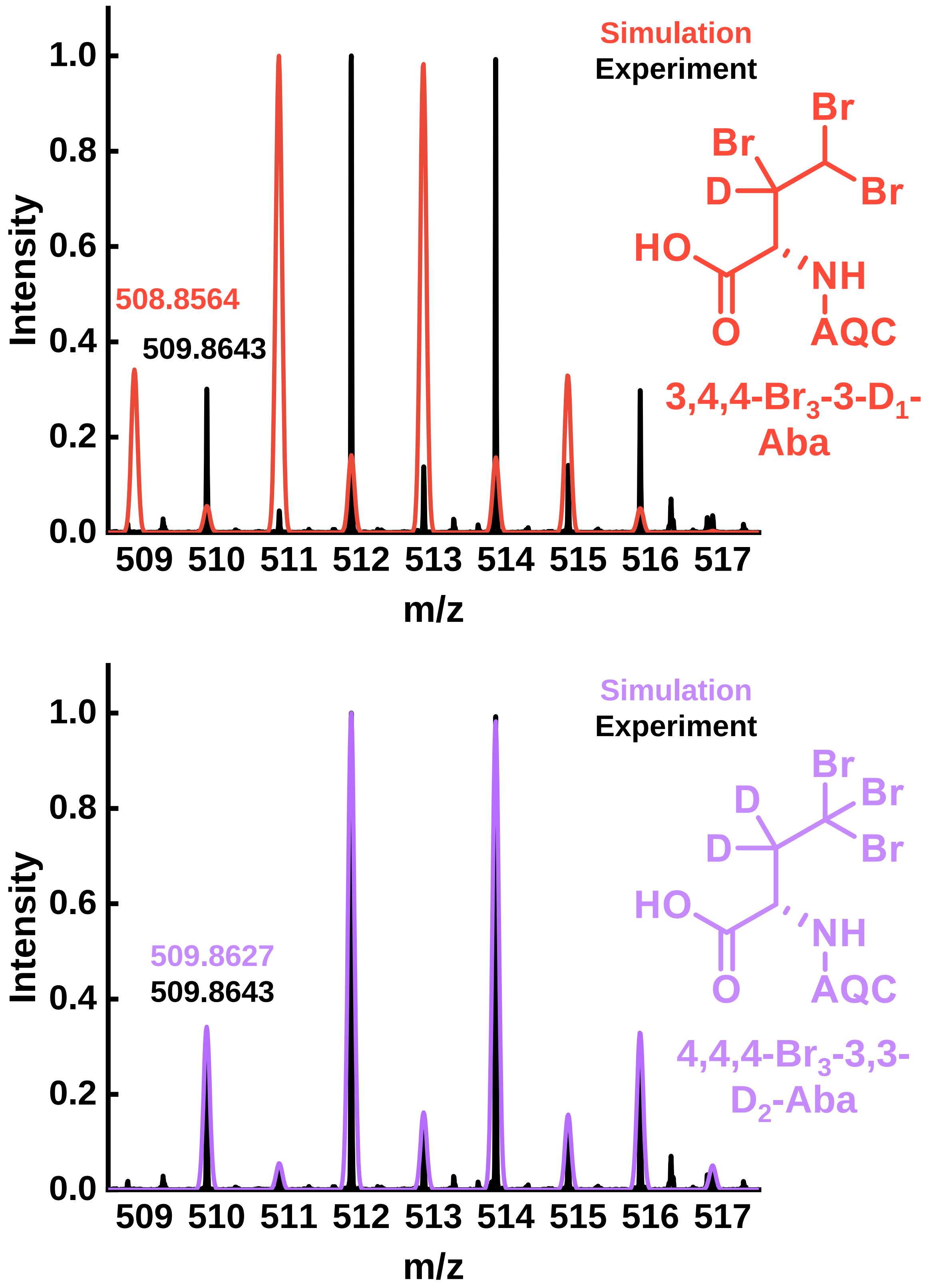


**Supplementary Figure 26.** Accurate mass spectra for the tri-brominated product with simulated 3,4,4-$Br_3$-3-$D_1$-Aba (top) and 4,4,4-$Br_3$-3,3-$D_2$-Aba (bottom). Isotope distribution pattern and experimental mass strongly agrees with the theoretical mass for 4,4,4-$Br_3$-3,3-$D_2$-Aba (monoisotopic mass error = 3.1 ppm), confirming that both deuterium atoms were retained in the bromination reaction, and all bromine atoms were installed at C4.

### Supplemental Info References:


1. Wilson, R. H.; Chatterjee, S.; Smithwick, E. R.; Dalluge, J. J.; Bhagi-Damodaran, A. Role of Secondary Coordination Sphere Residues in Halogenation Catalysis of Non-Heme Iron Enzymes. *ACS Catal.* **2022**, *12* (17), 10913–10924. https://doi.org/10.1021/acscatal.2c00954.

2. Mehmood, R.; Qi, H. W.; Steeves, A. H.; Kulik, H. J. The Protein's Role in Substrate Positioning and Reactivity for Biosynthetic Enzyme Complexes: The Case of SyrB2/SyrB1. *ACS Catal.* **2019**, *9* (6), 4930–4943. https://doi.org/10.1021/acscatal.9b00865

3. D.A. Case, K. Belfon, I.Y. Ben-Shalom, S.R. Brozell, D.S. Cerutti, T.E. Cheatham, III, V.W.D. Cruzeiro, T.A. Darden, R.E. Duke, G. Giambasu, M.K. Gilson, H. Gohlke, A.W. Goetz,R Harris, S. Izadi, S.A. Izmailov, K. Kasavajhala, A. Kovalenko, R. Krasny, T. Kurtzman, T.S. Lee, S. LeGrand, P. Li, C. Lin, J. Liu, T. Luchko, R. Luo, V. Man, K.M. Merz, Y. Miao, O. Mikhailovskii, G. Monard, H. Nguyen, A. Onufriev, F. Pan, S. Pantano, R. Qi, D.R. Roe, A. Roitberg, C. Sagui, S. Schott-Verdugo, J. Shen, C.L. Simmerling, N.R. Skrynnikov, J. Smith, J. Swails, R.C. Walker, J. Wang, L. Wilson, R.M. Wolf, X. Wu, Y. Xiong, Y. Xue, D.M. York and P.A. Kollman (2020), AMBER 2020, University of California, San Francisco.

4. Li, P.; Merz, K. M. Jr. MCPB.Py: A Python Based Metal Center Parameter Builder. *J. Chem. Inf. Model.* **2016**, *56* (4), 599–604. https://doi.org/10.1021/acs.jcim.5b00674

5. Gaussian 16, Revision C.01, M. J. Frisch, G. W. Trucks, H. B. Schlegel, G. E. Scuseria, M. A. Robb, J. R. Cheeseman, G. Scalmani, V. Barone, G. A. Petersson, H. Nakatsuji, X. Li, M. Caricato, A. V. Marenich, J. Bloino, B. G. Janesko, R. Gomperts, B. Mennucci, H. P. Hratchian, J. V. Ortiz, A. F. Izmaylov, J. L. Sonnenberg, D. Williams-Young, F. Ding, F. Lipparini, F. Egidi, J. Goings, B. Peng, A. Petrone, T. Henderson, D. Ranasinghe, V. G. Zakrzewski, J. Gao, N. Rega, G. Zheng, W. Liang, M. Hada, M. Ehara, K. Toyota, R. Fukuda, J. Hasegawa, M. Ishida, T. Nakajima, Y. Honda, O. Kitao, H. Nakai, T. Vreven, K. Throssell, J. A. Montgomery, Jr., J. E. Peralta, F. Ogliaro, M. J. Bearpark, J. J. Heyd, E. N. Brothers, K. N. Kudin, V. N. Staroverov, T. A. Keith, R. Kobayashi, J. Normand, K. Raghavachari, A. P. Rendell, J. C. Burant, S. S. Iyengar, J. Tomasi, M. Cossi, J. M. Millam, M. Klene, C. Adamo, R. Cammi, J. W. Ochterski, R. L. Martin, K. Morokuma, O. Farkas, J. B. Foresman, and D. J. Fox, Gaussian, Inc., Wallingford CT, 2016.

6. Tian, C.; Kasavajhala, K.; Belfon, K. A. A.; Raguette, L.; Huang, H.; Migues, A. N.; Bickel, J.; Wang, Y.; Pincay, J.; Wu, Q.; Simmerling, C. ff19SB: Amino-Acid-Specific Protein Backbone Parameters Trained against Quantum Mechanics Energy Surfaces in Solution. *J. Chem. Theory Comput.* **2020**, *16* (1), 528–552. https://doi.org/10.1021/acs.jctc.9b00591.

7. Wilson, R. H.; Zamfir, S.; Sumner, I. Molecular Dynamics Simulations Reveal a New Role for a Conserved Active Site Asparagine in a Ubiquitin-Conjugating Enzyme. *J. Mol. Graph. Model*. **2017**, 76, 403–411. https://doi.org/10.1016/j.jmgm.2017.07.006.

8. Roe, D. R.; Cheatham, T. E. PTRAJ and CPPTRAJ: Software for Processing and Analysis of Molecular Dynamics Trajectory Data. *J. Chem. Theory Comput.* **2013**, *9* (7), 3084–3095. https://doi.org/10.1021/ct400341p

9. Nicholls, A. Confidence Limits, Error Bars and Method Comparison in Molecular Modeling. Part 1: The Calculation of Confidence Intervals. *J. Comput. Aided Mol. Des*. **2014**, 28 (9), 887–918. https://doi.org/10.1007/s10822-014-9753-z.

10. Neese, F. Software Update: The ORCA Program System—Version 5.0. WIREs Computational Molecular Science 2022, 12 (5), e1606. https://doi.org/10.1002/wcms.1606.

11. Zhao, Y.; Truhlar, D. G. The M06 Suite of Density Functionals for Main Group Thermochemistry, Thermochemical Kinetics, Noncovalent Interactions, Excited States, and Transition Elements: Two New Functionals and Systematic Testing of Four M06-Class Functionals and 12 Other Functionals. Theor Chem Account 2008, 120 (1), 215–241. https://doi.org/10.1007/s00214-007-0310-x.

12. Weigend, F.; Ahlrichs, R. Balanced Basis Sets of Split Valence, Triple Zeta Valence and Quadruple Zeta Valence Quality for H to Rn: Design and Assessment of Accuracy. Phys. Chem. Chem. Phys. 2005, 7 (18), 3297–3305. https://doi.org/10.1039/B508541A.

13. Weigend, F. Accurate Coulomb-Fitting Basis Sets for H to Rn. Phys. Chem. Chem. Phys. 2006, 8 (9), 1057–1065. https://doi.org/10.1039/B515623H.

14. Cossi, M.; Rega, N.; Scalmani, G.; Barone, V. Energies, Structures, and Electronic Properties of Molecules in Solution with the C-PCM Solvation Model. Journal of Computational Chemistry 2003, 24 (6), 669–681. https://doi.org/10.1002/jcc.10189.

15. Armenta, J. M.; Cortes, D. F.; Pisciotta, J. M.; Shuman, J. L.; Blakeslee, K.; Rasoloson, D.; Ogunbiyi, O.; Sullivan, D. J. Jr.; Shulaev, V. Sensitive and Rapid Method for Amino Acid Quantitation in Malaria Biological Samples Using AccQ•Tag Ultra Performance Liquid Chromatography-Electrospray Ionization-MS/MS with Multiple Reaction Monitoring. Anal. Chem. 2010, 82 (2), 548–558. https://doi.org/10.1021/ac901790q.

16. Oss, M.; Kruve, A.; Herodes, K.; Leito, I. Electrospray Ionization Efficiency Scale of Organic Compounds. Anal. Chem. 2010, 82 (7), 2865–2872. https://doi.org/10.1021/ac902856t.
